# Effect of Biceps Brachii Muscle Stimulation on Respiration before and after Cervical Spinal Cord Injury

**DOI:** 10.64898/2025.12.30.697059

**Authors:** Natalia Shevtsova, Vitaliy Marchenko, Ilya Rybak, Tatiana Bezdudnaya

## Abstract

Cervical spinal cord injuries (SCI) often lead to respiratory impairments, significantly increasing morbidity and mortality in affected individuals. Limb muscle/afferent stimulation has been suggested as a potential approach to enhance breathing when supraspinal control over spinal respiratory circuits is compromised due to cervical SCI. Using a combination of intact, C2 Hemisected (C2Hx), and complete C1 Transected (C1Tx) rat models, we systematically evaluated the influence of forelimb muscle afferent input on phrenic motor output and respiratory patterns. Computational modeling was employed to replicate our experimental data and generate predictions. The developed computational model incorporates bilaterally located spinal and supraspinal respiratory circuits, allowing us to simulate their specific contributions to phrenic motor output under different conditions. In this study, we hypothesize that, in addition to supraspinal control, spinal circuits integrate limb sensory input to modulate the activity of phrenic motor neurons through local excitatory and inhibitory interneurons. These intraspinal pathways, normally suppressed by inhibition, can be recruited during movement to adapt breathing to motor demands. Our experimental and computational modeling results following biceps stimulation after C2Hx and C1Tx support this hypothesis, demonstrating that activation of limb afferent pathways can enhance phrenic motor output even after partial or complete loss of supraspinal drive. In the fully transected preparation, this effect required pharmacological disinhibition, confirming the presence of latent spinal pathways. This study provides the first evidence for functionally relevant intraspinal interactions between limb muscles and respiratory circuits and identifies a potential spinal mechanism that could be leveraged to promote breathing recovery after cervical SCI.

**Key Points:**

- Biceps brachii electrical stimulation can increase breathing frequency and tidal volume in spontaneously breathing rats and variably induce transient increases or entrainment of phrenic nerve activity in intact rats under controlled ventilation.
- Biceps stimulation enhances ipsilateral phrenic activity in acute C2 hemisected (C2Hx) rats.
- Biceps stimulation drives bilateral phrenic output in C1 transected (C1Tx) rats under conditions of pharmacologically induced spinal disinhibition.
- A computational model incorporating bilateral brainstem and cervical spinal respiratory circuits is developed to simulate cervical spinal cord injuries (C2Hx and C1Tx) and to examine how biceps stimulation affects respiratory activity under different conditions.
- These findings demonstrate a critical role for spinal circuits in locomotor–respiratory interactions.

## 1. Introduction

Breathing is a vital, continuous process that adapts dynamically gas exchange to environmental, metabolic, and behavioral demands. The neural circuits responsible for generating and regulating respiratory rhythm are distributed across multiple regions of the central nervous system. In mammals, the core respiratory rhythm originates from a central pattern generator (CPG) located in the lower brainstem, extending from the medulla to the pons (Richter, 1996; Smith et al., 2013; Smith et al., 2009). Within this network, the pre-Bötzinger complex (pre-BötC), situated in the ventrolateral medulla, plays a key role as a major source of rhythmic inspiratory activity (Koshiya & Smith, 1999; Rekling & Feldman, 1998; Smith et al., 2009; Smith et al., 1991). The Bötzinger complex (BötC), which primarily contains post-inspiratory and expiratory neurons (Ezure, 1990; Ezure et al., 2003; Khalilpour et al., 2024; Richter & Smith, 2014), works together with the pre-BötC to form the kernel of the respiratory CPG (Rybak et al., 2007; Smith et al., 2013). Oscillatory respiratory activity emerges from the intrinsic biophysical properties of neurons within these regions and their reciprocal synaptic interactions.

The activity of the CPG is modulated by multiple brainstem centers, including pontine nuclei, and transmitted through premotor neurons in the rostral and caudal ventral respiratory groups (rVRG and cVRG) to the spinal cord. Descending projections then activate phrenic, intercostal, and abdominal motoneurons, which drive the muscles required for respiration. Additional structures such as the retrotrapezoid nucleus (RTN)/parafacial respiratory group (pFRG) and the parabrachial (PB) and Kölliker-Fuse (KF) nuclei contribute to shaping respiratory output, particularly in coordinating phase transitions and generating active expiration(Smith, 2022).

Although brainstem mechanisms underlying respiratory rhythm generation have been studied extensively (Feldman et al., 2013; Feldman et al., 2003; Smith, 2022; Smith et al., 2013), the role of spinal circuits in respiratory control remains less understood. Emerging evidence suggests that spinal interneurons are essential for modulating respiratory activity both under physiological conditions and following spinal cord injury (SCI) (Jensen et al., 2019; Lane et al., 2008; Lin et al., 2025; Marchenko et al., 2015; Sandhu et al., 2009; Zholudeva et al., 2017). These interneurons integrate descending drive with sensory feedback, contributing to the adaptive regulation of breathing.

Interactions between locomotor and respiratory circuits represent another crucial dimension of respiratory control. During locomotion or exercise, sensory feedback from rhythmically moving limbs exerts a powerful influence on breathing. Two principal mechanisms, supraspinal and intraspinal coupling, have been proposed to explain these locomotor–respiratory interactions (Juvin et al., 2022; Shevtsova et al., 2019). While most research has focused on supraspinal mechanisms involving the brainstem, spinal pathways likely play an important role in shaping respiratory output through neuronal networks located within the spinal cord. Sensory afferents from limb muscles can project to both brainstem nuclei, and to local spinal interneurons that may directly influence phrenic motor output.

The present study aims to investigate the effect of electrical stimulation of the biceps brachii muscle on respiratory activity, focusing on phrenic nerve output before and after cervical spinal cord injury. By examining how forelimb muscle stimulation influences breathing under both intact and injured conditions, this work seeks to elucidate the spinal mechanisms underlying sensorimotor coupling in respiratory control and to explore potential pathways for promoting respiratory recovery after SCI.

## 2. Methods

### 2.1. Experimental methods

#### 2.1.1. General surgical procedures and animals

All surgical and experimental procedures were approved by the Drexel University Institutional Animal Care and Use Committee and adhered to AAALAC/NIH guidelines. All experiments were performed on adult Sprague-Dawley rats (360-420g) of both sexes. Animals were initially anesthetized with Isoflurane (2% in 100% O2) through the mask. Animal was placed on servocontrolled heating blanket coupled to a rectal thermometer (Harvard Apparatus) to keep the temperature at 37.0 C during surgery and all recordings. Plastic cannula was inserted into the trachea following tracheotomy, and animal was connected to artificial ventilation (Columbus Apparatus) while maintaining the same level of anesthesia. The femoral vein and artery were catheterized for drug delivery and measuring blood pressure, respectively. Both phrenic nerves were gently dissected from surrounding tissue and cut at their distal ends. Chemoreceptor and baroreceptor denervation was achieved by cutting the carotid sinus nerves bilaterally and sectioning the aortic depressor nerves, effectively eliminating peripheral chemoreceptor and baroreceptor inputs. Vagotomy was performed by cutting two vagus nerves. The internal carotid arteries were ligated bilaterally to minimize bleeding during decerebration. Animals transected at C1 cervical level were maintained under isoflurane anesthesia and were not decerebrated. Recordings in all animals were performed after 3-4 hours post-C2Hx or C1Tx.

#### 2.1.2. Experiments on spinal cord intact rats

Two groups of animals were used for experiments on spinal cord intact rats. First group: we used spontaneously breathing, decerebrate rats (n = 6). The animals were maintained under a low dose of isoflurane (∼1.0%) to slightly immobilize them, as biceps stimulation could otherwise produce movements or escape reactions. Oxygen was delivered continuously via a tracheal T-cannula at low pressure, where expired air directed to a capnograph and airflow meter. This setup allowed the animals to maintain autonomous breathing, while tidal CO₂ levels and airflow were continuously monitored. Parameters of stimulation: closed-loop stimulation triggered by expiratory-phase tracheal pressure; pulse duration, 0.5 ms; burst frequency, 100–300 Hz; number of pulses per burst, 10–25; stimulation intensity, 0.5–2 mA (3-5T).

Second group of animals: We used phrenic nerve (PN) recording in decerebrate, vagotomized, and chemo– and baroreceptor-denervated rats (n = 12). These animals were maintained under continuous infusion of vecuronium bromide (3–4 mg/kg/h) dissolved (0.5 mg/ml) in Ringer-Locke solution to immobilize them and eliminate all respiratory artifacts during recordings. Chemo– and baro-denervation was performed to eliminate any other peripheral influence on biceps stimulation effects on respiration. The following stimulation parameters were used: pulse duration 0.5–1 ms, burst frequency 50–100 Hz, number of pulses 15–30, intensity 0.2–1 mA, and inter-burst interval 1.1–1.5 s.

#### 2.1.3. Experiments on C2Hx rats

All C2Hx animals were decerebrate, vagotomozed and chemobarodenervated (n=6). The decerebration procedure has been described in detail previously (Bezdudnaya et al., 2017). Briefly, for decerebration, artificially ventilated (isoflurane 2%) rat was placed into a stereotaxic frame, and dexamethasone (1mg/kg, i.m.) was injected to prevent brain edema. Arterial blood pressure was continuously monitored. After biparietal craniotomies, the superior sagittal sinus was ligated, and the brain axes was transected at the rostral border of the superior colliculus. Cerebral tissue rostral to the transection was aspirated, and small pieces of gelfoam soaked with a cold thrombin solution were placed into the skull cavity to stop bleeding. The rats were allowed to recover for 3-4 hours after decerebration. During this time, 1ml of dextrose (5%) was injected intravenously (every 30 min) to stabilize blood pressure. Subsequently, first seven spinal segments of the spinal cord were surgically exposed. The animal was centered within the stereotactic frame, and the vertebral column was raised using the T2 vertebral spinous process to align the spinal cord. A bilateral pneumothorax was performed to eliminate artifacts associated with respiratory movement during electrophysiological recordings. Anesthesia was withdrawn and a paralytic agent was continuously injected (vecuronium bromide; 3-4 mg/kg/h, 0.5 mg diluted in 1 ml Ringer-Loker’s solution). Once an animal stabilized post-decerebration, the lateral hemisection was made under 2% isoflurane anesthesia with a scalpel blade (11 size) immediately below the C2 spinal rootlets (Bezdudnaya et al., 2017). Anesthesia was slowly withdrawn during 1 h and all recordings were performed started 3 h post-C2Hx.

Stimulations were performed via two insulated tungsten wires inserted within the muscle. Closed-loop stimulation was employed. Stimulation was triggered by contralateral phrenic activity in the beginning of respiration (Fig. 1A). The following stimulation parameters were used: pulse duration 1 ms, burst frequency 50–100 Hz, number of pulses 5–10, and intensity 0.6–1.5 mA (2–5T).

**Figure. 1.**
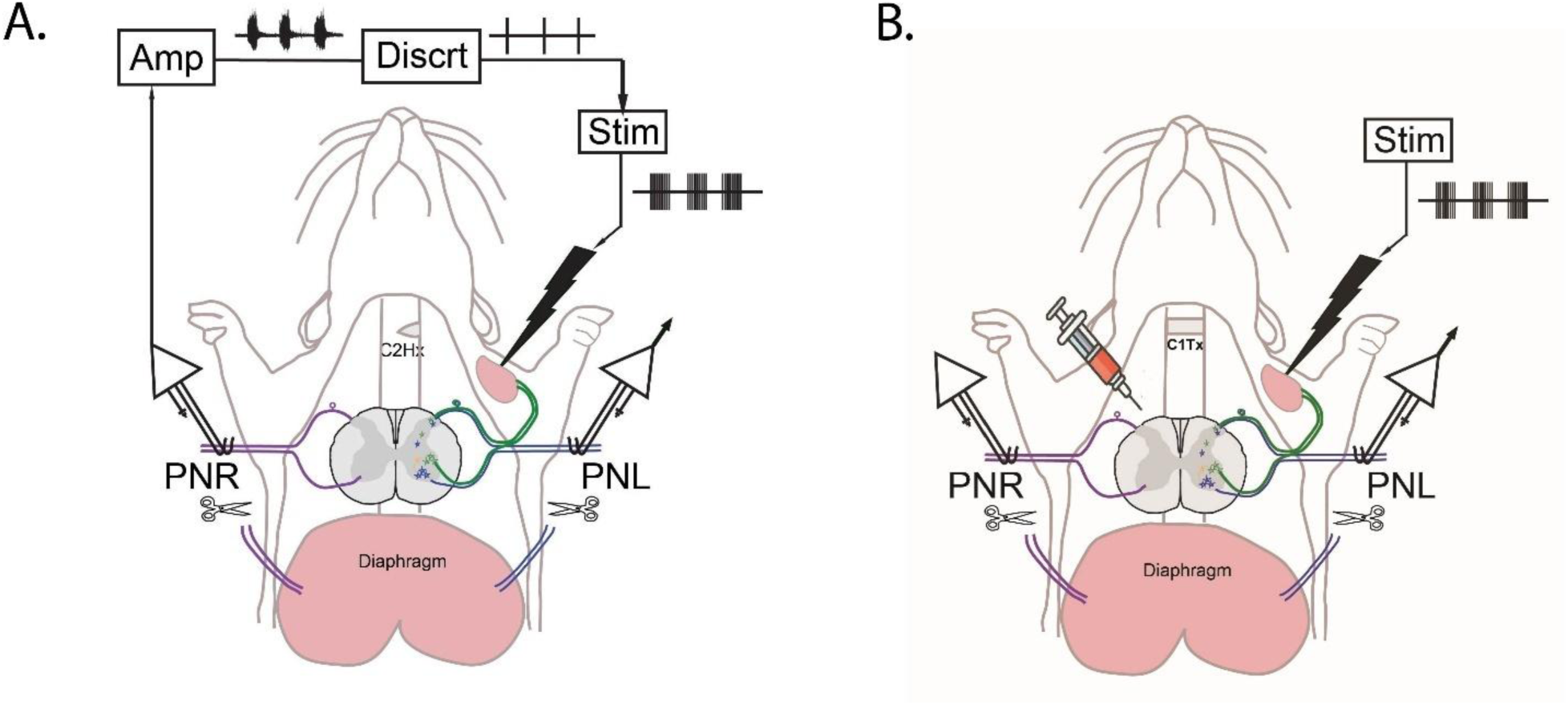
A. Schematic of biceps stimulation experiment is C2Hx rats. B. Schematic of unilateral biceps stimulation experiment in C1Tx rats. PNR – right phrenic nerve, PNL – left phrenic nerve, Amp-Amplifier, Discrt – Discriminator, Stim – Stimulator. Syringe indicates drug application to the spinal cord.

#### 2.1.4. Experiments on C1Tx rats

Complete C1 transection was performed below the C1 spinal rootlets (Bezdudnaya, Lane, et al., 2018; Bezdudnaya et al., 2020). Following transection, the rats were allowed to recover for several hours before recording. After 4 hours of C1Tx cocktail containing 100-200 μMol Gabazine (GABAz)+Strychnine (STR) were applied over C3-C5 cervical segments. We recorded both phrenic activity in response to unilateral (n=9) and bilateral (n=6) biceps electrical stimulation. C1Tx animals were recorded under isoflurane anesthesia (1.5%). Open-loop biceps stimulation was employed. The following stimulation parameters were used: stimulation frequency 0.1 Hz, pulse duration 0.5 ms, burst frequency 100 Hz, number of pulses 3, and intensity 0.7–1.4 mA.

#### 2.1.4. Electrophysiological recordings

All electrophysiological signals were digitized and recorded on PC using Chart5 software (AD Instruments). All recordings were performed during eupneic ventilation (O2 50%, end-tidal CO2 – 4.5-5%). Bilateral phrenic nerve recordings were performed in mechanically ventilated and immobilized rats. A mineral oil bath was created on both sides of the spinal cord to immerse nerves and bipolar silver electrodes for nerve recording. Nerve activity were amplified (1000x) and filtered (10-5,000 Hz) using differential amplifier Neurolog (Digitizer, UK). End tidal CO2 level, arterial blood pressure, and trachealpressure were continuesly recorded. Intrathecal drug delivery were performed as described before (Bezdudnaya et al., 2020).

#### 2.1.5. Experimental data analysis

All data were converted to ASCII tables and analyzed using Matlab and SigmaPlot 11. Mean values of tidal volume (Tv), amplitude of integrated phrenic nerve (PN) activity (integration time constant τ = 0.05 s), inspiratory and expiratory phase durations (Ti and Te), and respiratory frequency were calculated using custom made scripts. Normality was assessed with the Shapiro–Wilk test. Pairwise comparisons of respiratory parameters were performed using paired t-tests or, when appropriate, the nonparametric Wilcoxon rank-sum test. Data are presented as mean ± standard error (M ± SE). Statistical significance was defined at *p* < 0.05.

### 2.2. Computational modeling

#### 2.2.1. Simulation tools

Computational modeling studies were performed using the neural simulation package NSM 2.5.7 developed in the Computational Neuroscience group at Drexel University. Differential equations were solved using the exponential Euler integration method with a step size of 0.1 ms. Previously, it was used for many published, well-known models of neural control of respiration (Molkov et al., 2010; Molkov et al., 2017; Molkov et al., 2014; Potts et al., 2005; Rybak, Shevtsova, Paton, et al., 2004; Shevtsova et al., 2014; Shevtsova et al., 2011; Smith et al., 2013; Smith et al., 2007) and locomotion (Danner et al., 2019; Rybak et al., 2015; Rybak et al., 2025; Rybak et al., 2013; Rybak, Shevtsova, et al., 2006; Rybak et al., 2024; Rybak, Stecina, et al., 2006; Shevtsova, 2013; Shevtsova et al., 2022; Shevtsova NA, 2016; Shevtsova et al., 2015; Zhang et al., 2022; Zhong et al., 2012)

The package uses parallel computing to increase the number of populations and neurons per population with more extensive randomization of neuronal parameters within their physiological ranges to approximate biological reality and provide a more stable model behavior.

#### 2.2.2. Modeling approach

The model represents a bilateral symmetric network of interacting neural populations and includes the left and right brainstem respiratory circuits and the left and right spinal compartments. The left and right brainstem cord respiratory circuits were built upon our previous computational models of the brainstem respiratory network (Molkov et al., 2010; Potts et al., 2005; Rybak, Shevtsova, Paton, et al., 2004; Smith et al., 2007) and incorporated the medullary (MEDULLA) and pontine (PONS) compartments. Each of these compartments included multiple neuron population (see Results). For simplicity and similar to the previous modeling studies, we assumed that each compartment contains only populations of respiratory neuron types that are known to be dominantly present in this compartment. The simplified left and right spinal compartments included populations of phrenic motoneurons (PhMN), simplified models of lungs and slowly adapting pulmonary stretch receptors, and sensory interneurons populations (InS).

##### Modeling single neurons

All neurons were modeled in the Hodgkin-Huxley style (single-compartment models) and incorporated known biophysical properties and channel kinetics characterized in respiratory neurons in vitro. Specifically, the kinetics of the fast sodium and the persistent (slowly inactivating) sodium channels were described using the experimental data obtained in studies of neurons from the rat RVLM (Rybak, Ptak, et al., 2003); the kinetics of high-voltage-activated calcium current was described based on the study of calcium currents in rat medullary neurons in vitro (Elsen & Ramirez, 1998); the intracellular calcium dynamics were described using data by Frermann et al. (Frermann et al., 1999). The descriptions of other ion channels, e.g., the potassium rectifier and calcium-dependent potassium ones, and all other cellular parameters were derived from our previous models (Rybak et al., 1997a, 1997b; Rybak, Shevtsova, Paton, et al., 2004; Rybak, Shevtsova, Ptak, et al., 2004; Rybak, Shevtsova, et al., 2003). A full description of single neuron model, description of channel kinetics, and major model parameters can be found in Appendix (or Supplemental. Materials).

##### Modeling neural populations

The important advantage of the NSM package is that it allows a user to explicitly construct neural models at the level of populations, yet having each neuron modeled using Hodgkin-Huxley formalisms. A user defines the average values of neuron parameters and their variances, as well as the probabilities of synaptic connection and average values and variances of connection weights between neuron populations. Then, parameters of individual neurons and connections are assigned, using a random generator. Parameter randomization provides robust model behavior and allows realistic simulation of neuron populations.

In the present model, each functional type of neuron was represented by a population of 20-100 neurons. The heterogeneity of neurons within each population was set by a random distribution of the reversal potentials of the leakage channels, *E_L_* (mean values ± SD, SD was chosen to be equal 2% of mean value) and initial conditions for values of membrane potential, calcium concentrations and channel conductances.

Connections between the populations were established so that, if a population A was assigned to receive an excitatory or inhibitory input from a population B or external drive D, then each neuron of population A received the corresponding excitatory or inhibitory synaptic input from each neuron of population B or from drive D, respectively. In all simulations, a settling period of 20 s was allowed in each simulation before data were collected. Each simulation was repeated 20-30 times and demonstrated qualitatively similar behaviour for particular values of the standard deviation of *E_L_* and other distributed parameters.

##### Stimulating effects challenging experimental conditions and afferent stimulation

The NSM package has special tools for simulating various *in vivo* and *in vitro* experimental approaches, including lesions or transections, different challenging experimental conditions, as well as simulation of various stimulation types. The effect of vagotomy was simulated by removal of the feedback from lungs to the respiratory compartment of the model. To simulate, the effect of hypercapnia we used the same approach as in our previous modeling studies (Molkov et al., 2010; Molkov et al., 2014) and applied additional external drive to the late-E population located in the RTN/pFRG compartment (for detail, see **Modeling Results**). To simulate the effect of hypoxia we used a novel approach and applied an additional excitatory drive to the Ine-NTS population, located in the NTS compartment. The effect of stimulation of muscle afferents was simulated by application of a periodic external excitatory input to the sensory interneuron population, InS (see **Modeling Results**). During these experiments we monitored simulated phrenic and hypoglossal activities and compared them to the corresponding experimental results. Simulated phrenic and hypoglossal activities were represented as average activities of neurons (spikes/per neuron/per second) in the PhMn and HgMn populations, respectively. In the current study, we simulated the ipsi– and contralateral effects of biceps stimulation on respiratory pattern.

To simulate a partial or complete elimination of supraspinal control in C2Hx or C1Tx conditions (Fig. 4-5) removed half or all connections between the brainstem and spinal compartments in the model by setting the corresponding connection weights to zero (see Fig. M1).

## 3. Results

### 3.1. Experimental results

#### 3.1.1. Effects of biceps stimulation in intact rats

All experiments reported in this chapter were performed in decerebrate, spinal cord intact animals.

In the first set of experiments, we used spontaneously breathing rats and recorded changes in airflow before and during biceps brachii muscle stimulation (n=6). Biceps stimulation had two effects: 1) an increase in tidal volume (2.8±0.4 vs 3.4±0.13 ml; p=0.008) and 2) an increase in breathing frequency (90.2± 7.9 vs 105.01± 6.3 breaths/min; p=0.002) (Figure 2A and B). However, these effects decayed quickly.

**Figure 2.**
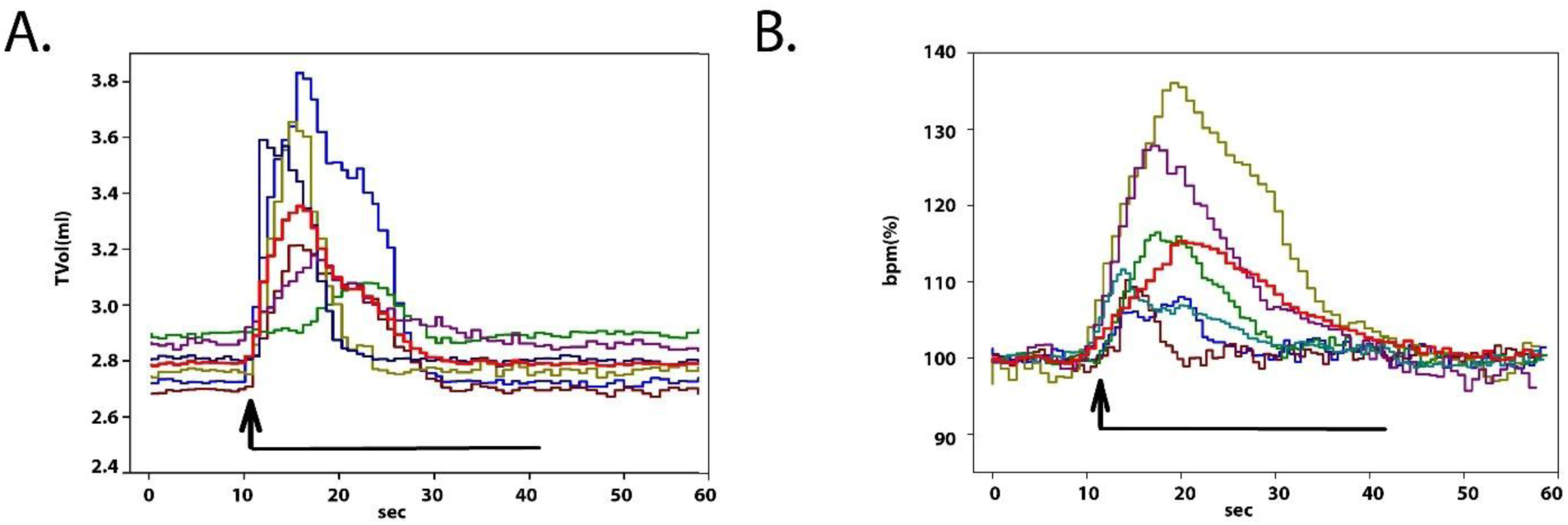
Effect of biceps stimulation on tidal volume and frequency of breathing in spontaneously breathing decerebrate rats. A. Changes in tidal volume during biceps stimulation (n = 6). B. Changes in breathing frequency during biceps stimulation (n = 6). Red line indicates averaged traces. TV – tidal volume; BPM – breaths per minute. BPM values were normalized to pre-stimulation.

In the second set of experiments, we used vagotomized, chemo– and baro-denervated, mechanically ventilated rats (n=12). Phrenic nerve activity was recorded before and during biceps stimulation. These experiments produced inconsistent results, with two distinct responses observed: (1) a transient (duration: 2.8± 0.97 sec) increase in phrenic frequency, from 43.05±2.5 to 59.5±3.5 breaths/min (p<0.001; n=7, 58%); or (2) entrainment of phrenic nerve activity by biceps brachii stimulation (n=5, 42%) Although phrenic burst amplitude showed a slight increase during stimulation, these changes were not statistically significant (transient response: n = 7, 0.038 ± 0.02 vs. 0.041 ± 0.03 au, p = 0.052; entrainment: n = 5, 0.040 ± 0.003 vs. 0.042 ± 0.004 au, p =0.166). Representative examples of both transient frequency increases and entrainment are shown in Fig. 3.

**Figure 3.**
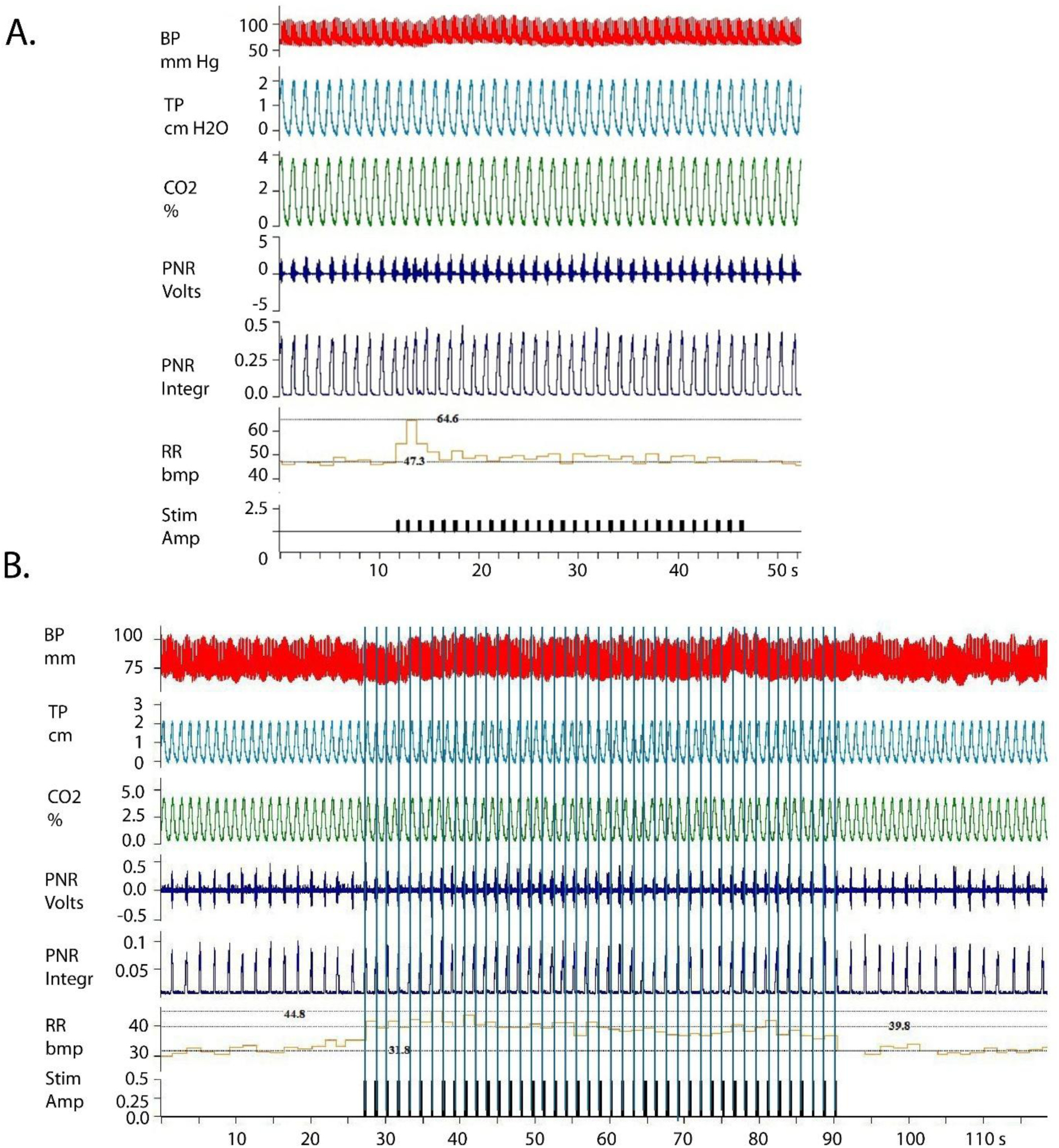
Representative traces showing the effect of biceps stimulation on phrenic nerve activity in intact rats. A. Example of transient increase of phrenic nerve frequency by biceps stimulation. B. Example of phrenic nerve entrainment by biceps stimulation. Abbreviations: BP-blood pressure, TP –tracheal pressure, CO2 –end tidal CO2, PNR-right phrenic activity, PNR-Intg-integrated phrenic activity, RR-respiratory rate, Stim – biceps stimulation.

#### 3.1.2. Effects of biceps stimulation in C2Hx rats

Electrical stimulation of the biceps brachii muscle was used to assess the effect of forelimb afferent inputs on phrenic motoneurons, spinal interneurons, and respiration (n=6). Closed-loop biceps stimulation ipsilateral to the C2Hx was delivered at the onset of each breath. Our experiments demonstrated a significant increase in phrenic nerve amplitude on the side ipsilateral to the injury compared to pre-stimulation values (p = 0.015). Interestingly, contralateral phrenic activity was not significantly affected during stimulation (p =0.105). A representative example of phrenic nerve activity before and during stimulation is shown in Figure 4A. Quantifications of phrenic nerve amplitudes ipsilateral and contralateral to the injury are shown in Figure 4B.

**Figure 4.**
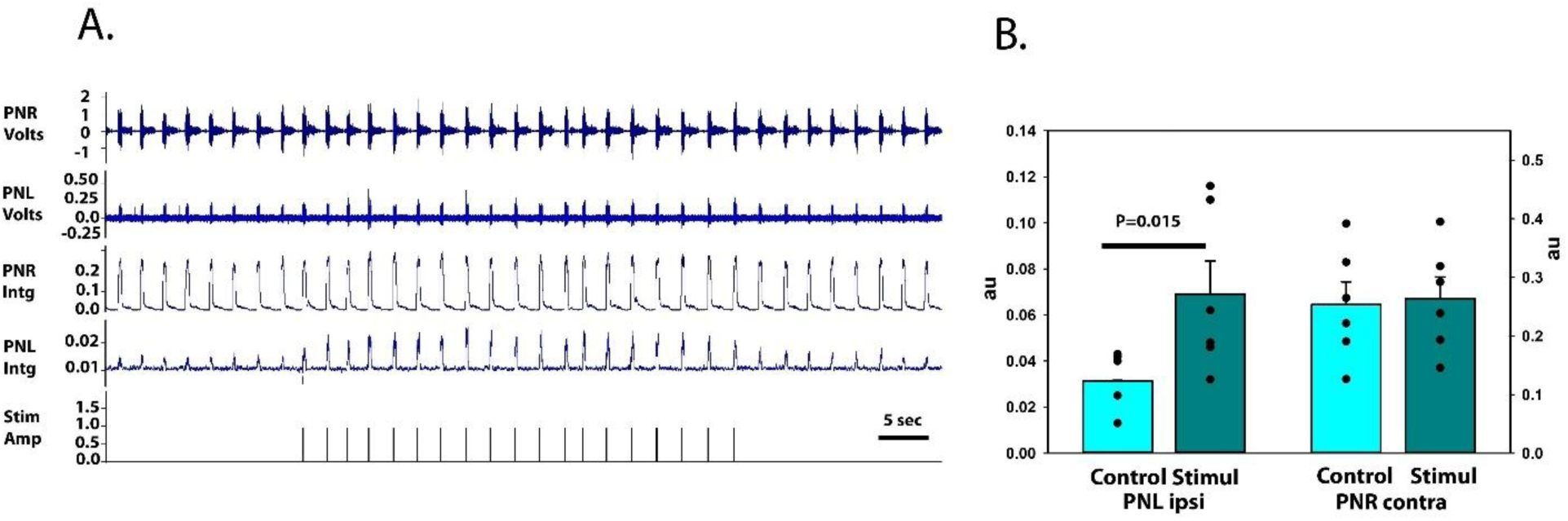
Effect of biceps stimulation in C2Hx rats. A. Example of biceps stimulation on phrenic nerve activity in C2Hx rat. B. Changes in phrenic amplitude during closed loop biceps stimulation in ipsilateral (left) and contralateral (right) nerves. PNL – left phrenic nerve, PNR – right phrenic nerve, Intg-integrated nerve activity (n=6).

#### 3.1.3. Effects of biceps stimulation in C1Tx rats

C1 transection (C1Tx) eliminated all descending influences from the brainstem to the phrenic spinal circuits, resulting in a total loss of phrenic nerve activity. Electrical stimulation of the biceps muscle performed several hours after C1Tx had no effect on phrenic nerve activity (data not shown). To investigate further, we applied a drug-induced disinhibition protocol using a cocktail of Gabazine and Strychnine. Fifteen minutes after drug application, the biceps brachii muscle was stimulated unilaterally and bilaterally. Each stimulation evoked bilateral phrenic bursts, with amplitudes significantly higher on the ipsilateral side of stimulation (p = 0.0161) and shorter latencies (24±3 ms) compared to the contralateral side (41 ± 3 ms, p = 0.0229). When biceps stimulation was applied bilaterally, the evoked phrenic burst amplitudes and latencies did not differ significantly between sides (p=0.5 and p=0.76 respectively). A representative example is shown in Figure 5A–B, with quantification presented in Figure 5C–D.

**Figure 5.**
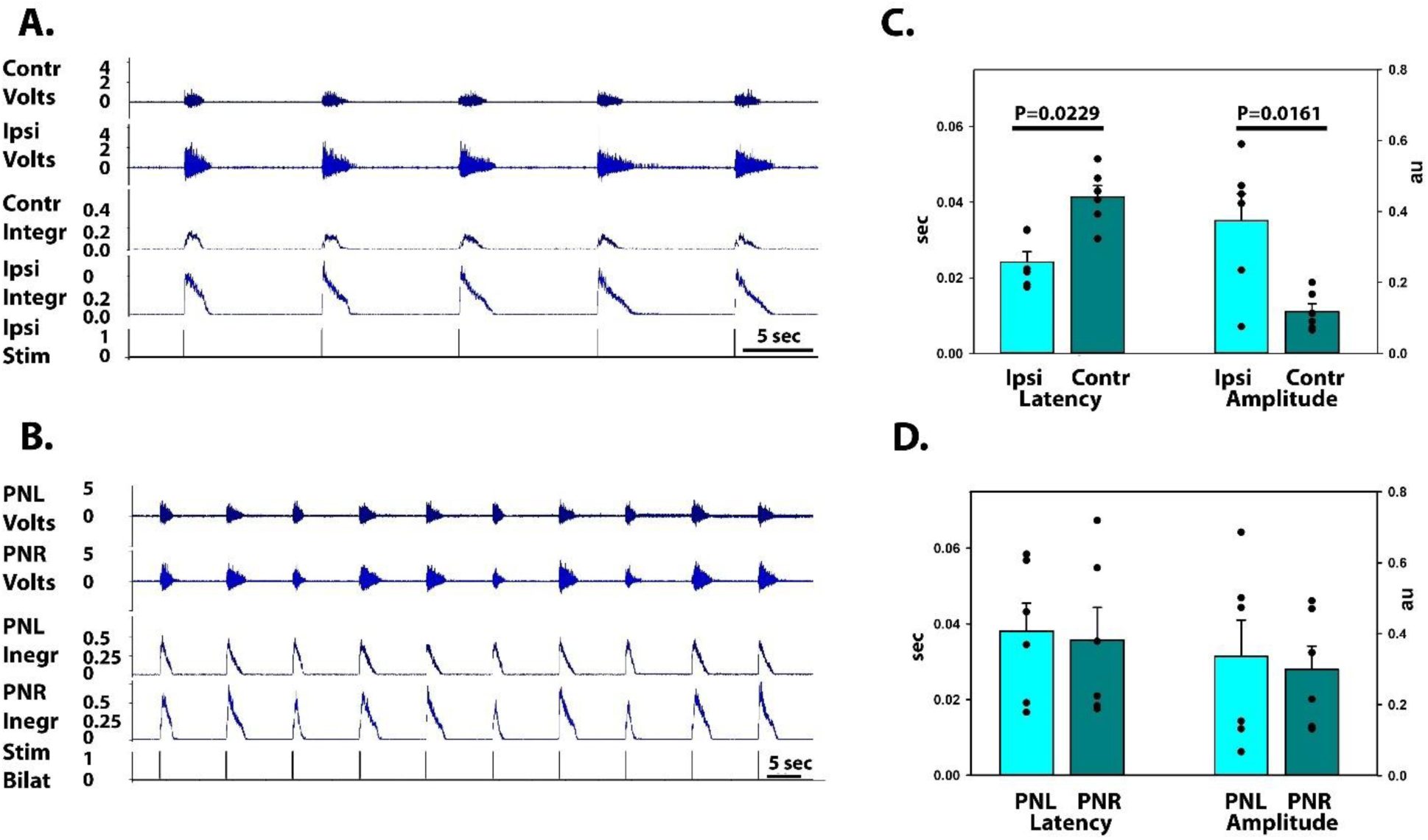
Effect of biceps stimulation in C1Tx rats after pharmacological disinhibition. A. Example of unilateral biceps stimulation; B. Example of bilateral biceps stimulation. C. Ipsi– and contra-lateral phrenic nerve latencies and amplitudes to unilateral biceps stimulation (n=6); D. Ipsi– and contra-lateral phrenic nerve latencies and amplitudes to bilateral biceps stimulation (n=6). PNL – left phrenic nerve, PNR – right phrenic nerve, Intg-integrated nerve activity.

### 3.2. Computational modeling

#### 3.2.1. Model structure

Most of existing computational models of the respiratory network focused primarily on brainstem circuits and described unilateral circuits, without accounting for interactions between the left and right respiratory centers. In addition to their functional operations, such bilateral interactions may play an important role in recovery of respiratory function after unilateral spinal cord injury. To address this issue, we developed a computational model of bilateral brainstem–spinal cord respiratory circuits that incorporates both left and right brainstem and spinal cord compartments. The model schematic is shown in Fig. M1. The left and right brainstem cord respiratory circuits incorporated the medullary (MEDULLA) and pontine (PONS) compartments. The core rhythmogenic components in MEDULLA included the Bötzinger (BötC) and pre Bötzinger (pre-BötC) complexes and rostral (rVRG) and caudal (cVRG) ventral respiratory groups, as well as other main respiratory compartments (raphé nucleus, retrotrapezoid nucleus/parafacial respiratory group (RTN/pFRG), and nucleus tractus solitaris (NTS), see Fig. M1). The detailed description of structure of and interactions within and between respiratory populations in and between the brainstem compartments in the model is given elsewhere (Molkov et al., 2010; Potts et al., 2005; Rybak, Shevtsova, Paton, et al., 2004; Smith et al., 2007) and only a short summary is provided here.

Specifically in our model, the BötC contains two populations of inhibitory expiratory neurons, the augmenting expiratory (aug-E) and the post-inspiratory (post-I) neurons, inhibiting neural populations within the pre-BötC and rVRG and each other during expiration (Fig. M1). In addition, he BötC contained an excitatory population (post-I(e)) that contributed to the post-I component of motor output of the central vagus (VN) nerve. All BötC neurons (comprising the post-I, post-I(e), and aug-E populations) had intrinsic adapting properties defined by the high-voltage activated calcium (*I*_CaL_) and calcium-dependent potassium (*I*_KCa_) currents in these neurons (see Methods and Tables 1 and 2).

**Figure M1.**
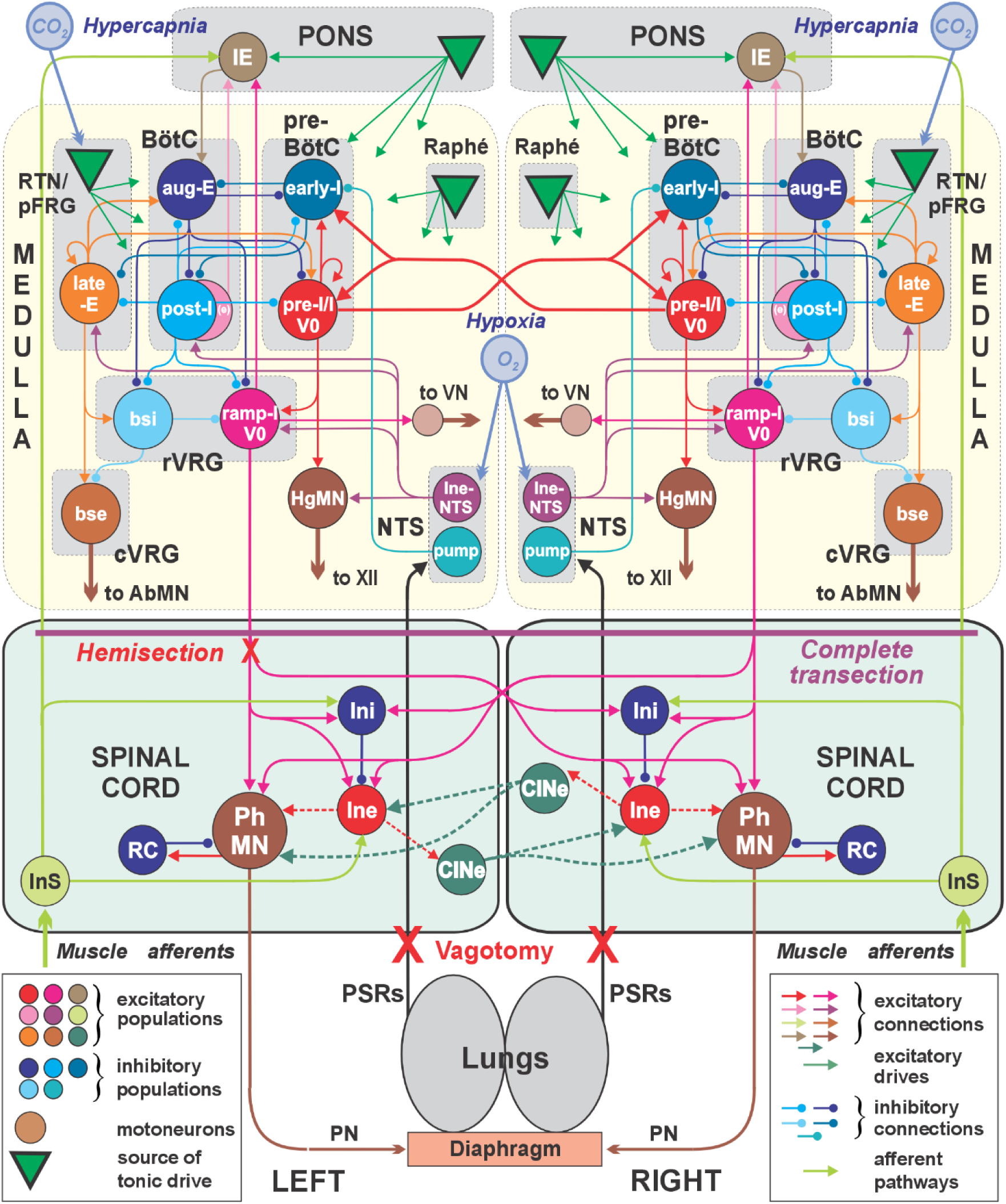
Schematic of the bilateral model of brainstem-spinal cord respiratory circuits. The model includes the left and right respiratory neural networks, each incorporating the pontine (PONS), medullary (MEDULLA) and spinal (SPINAL CORD) compartments. MEDULLA includes the BötC, pre-BötC, and rVRG compartments and the RTN, Raphe, and NTS nuclei. PONS includes the IE populations with inspiratory-expiratory modulated activity. SPINAL CORD includes the PhMNs and Renshaw cells (RC) recurrently inhibiting the PhMNs, and populations of local respiratory interneurons. Populations of sensory neurons, InS, mediate simulated limb afferent feedback (light green lines) to the pontine IE population and spinal respiratory excitatory interneurons (Ine), mediating afferent and supraspinal inputs to the PhMNs. The left and right networks will interact via V0 neurons at the level of the medulla and by commissural (CINe) populations at the spinal level. Model motor outputs are presented by simulated activities of the phrenic (PN), hypoglossal (XII), vagus (VN) and abdominal (AbN) nerves (thick red lines). The model includes a simplified lung model. Each population contains 20-100 Hodgkin-Huxley neurons.

**Table 1.** Steady state activation and inactivation variables and time constants for voltage-dependent ionic channels.

| Ionic channels | $m_{\infty}(V)$, $V$ in mV; $\tau_m(V)$, ms; $h_{\infty}(V)$, $V$ in mV; $\tau_h(V)$, ms |
| --- | --- |
| Fast sodium,<br>$Na$ | $m_{\infty Na} = 1/(1 + \exp(-(V + 43.8)/6))$;<br>$\tau_{mNa} = \tau_{mNa \max} / \cosh((V + 43.8)/14)$, $\tau_{mNa \max} = 0.252$;<br>$h_{\infty Na} = 1/(1 + \exp((V + 67.5)/10.8))$;<br>$\tau_{hNa} = \tau_{hNa \max} / \cosh((V + 67.5)/12.8)$, $\tau_{hNa \max} = 8.456$. |
| Persistent sodium,<br>$NaP$ | $m_{\infty NaP} = 1/(1 + \exp(-(V + 47.1)/3.1))$;<br>$\tau_{mNaP} = \tau_{mNaP \max} / \cosh((V + 47.1)/6.2)$, $\tau_{mNaP \max} = 1$;<br>$h_{\infty NaP} = 1/(1 + \exp((V + 60)/9))$;<br>$\tau_{hNaP} = \tau_{hNaP \max} / \cosh((V + 60)/9)$, $\tau_{hNaP \max} = 4000$. |
| Delayed rectifier<br>potassium,<br>$K$ | $\alpha_{\infty K} = 0.01 \cdot (V + 44)/(1 - \exp(-(V + 44)/5))$;<br>$\beta_{\infty K} = 0.17 \cdot \exp(-(V + 49)/40)$;<br>$m_{\infty K} = \alpha_{\infty K}/(\alpha_{\infty K} + \beta_{\infty K})$;<br>$\tau_{mK} = \tau_{mK \max}/(\alpha_{\infty K} + \beta_{\infty K})$; |
| High- voltage activated<br>calcium,<br>$Ca_L$ | $m_{\infty CaL} = 1/(1 + \exp(-(V + 27.4)/5.7))$;<br>$\tau_{mCaL} = 0.5$;<br>$h_{\infty CaL} = (1 + \exp((V + 52.4)/5.2))$;<br>$\tau_{hCaL} = 18$. |

**Table 2.** Maximal conductances of ionic channels in different neuron types.

| Neuron population<br>(location) | $\bar{g}_{Na}$, nS | $\bar{g}_{NaP}$, nS | $\bar{g}_K$, nS | $\bar{g}_{CaL}$, nS | $\bar{g}_{KCa}$, nS | $g_L$, nS | $E_L$, mV |
| --- | --- | --- | --- | --- | --- | --- | --- |
| IE (PONS) | 400 |  | 250 |  |  | 6 | -60 |
| pre-I /I (pre-BötC) | 170 | 5 | 180 |  |  | 2.5 | -68 |
| early-I (pre-BötC) | 400 |  | 250 | 0.005 | 10 | 6 | -60 |
| post-I (BötC) | 400 |  | 250 | 0.005 | 10 | 6 | -60 |
| post-I (e) (BötC) | 400 |  | 250 | 0.005 | 10 | 6 | -60 |
| aug-E (BötC) | 400 |  | 250 | 0.005 | 10 | 6 | -60 |
| late-E (RTN/pFRG) | 170 | 5 | 180 |  |  | 2.5 | -68 |
| ramp-I (rVRG) | 400 |  | 250 |  |  | 6 | -60 |
| bsi (rVRG) | 400 |  | 250 |  |  | 6 | -60 |
| bse (cVRG) | 400 |  | 250 |  |  | 6 | -60 |
| lne-NTS (NTS) | 400 |  | 250 |  |  | 6 | -60 |
| pump (NTS) | 400 |  | 250 |  |  | 6 | -60 |
| HgMN | 400 |  | 250 |  |  | 6 | -60 |
| Ini (SC) | 400 |  | 250 |  |  | 6 | -60 |
| lne (SC) | 400 |  | 250 |  |  | 6 | -60 |
| CINe (SC) | 400 |  | 250 |  |  | 6 | -60 |
| RhMN (SC) | 400 |  | 250 |  |  | 6 | -60 |
| RC (SC) | 400 |  | 250 |  |  | 6 | -60 |
| InS (SC) | 400 |  | 250 |  |  | 6 | -60 |

The pre-BötC includes two neural populations: excitatory pre-I/I and inhibitory early-I(1) (Fig. M1). The pre-I/I population serves as a major source of inspiratory activity in the model and projects to the ramp-I population of premotor inspiratory neurons of rVRG and, through a hypoglossal motoneuron population (HgMN), to the hypoglossal motor output (HN). The pre-I population comprises excitatory neurons with *I*_NaP_-dependent endogenous bursting properties and mutual excitatory synaptic connections within the population (see Appendix and Smith et al. (2007)). In our model (under normal conditions), most pre-I/I neurons operate in a tonic-spiking mode and is inhibited by expiratory neurons (post-I and aug-E) during expiration (for detail, see (Smith et al., 2007)). The early-I(1) population of pre-BötC is considered as a population of inhibitory interneurons with adapting properties (defined by *I*_CaL_ and *I*_KCa_, see Methods and Tables 1 and 2 in Appendix). This population receives excitation from the pre-I population and serves as a major source of inspiratory inhibition (Smith et al., 2007). This population inhibits all expiratory neurons during inspiration (Fig. M1). The rhythmic respiratory pattern (alternating inspiratory and expiratory phases) emerges from dynamic interactions between the pre-I/I and early-I populations located in the pre-BötC (active during inspiration) and the post-I and aug-E populations located in the BötC (active during expiration).

The rVRG compartment includes the excitatory ramp-I population and the inhibitory bulbospinal premotor (bsi) population (Fig. M1). Ramp-I is a population of excitatory premotor inspiratory neurons projecting to population of phrenic motoneurons (PhMN) driving the diaphragm. Activity of this population defines the phrenic motor output (PN) and the inspiratory component of the vagus nerve (VN) discharge. The major role of the inhibitory bsi population (with adapting neurons containing *I*_CaL_ and *I*_KCa_, see Appendix) is to shape the augmenting patterns of ramp-I neurons (Smith et al., 2007). The medullary populations received tonic excitatory drives from the raphè, RTN/pFRG, and pons.

The original Smith et al. (2007) model has been extended by incorporating the pontine IE population, the late-E population in the RTN/pFRG compartment (as in (Molkov et al., 2010)), the cVRG compartment comprising the excitatory bulbospinal premotor interneurons (bse), and populations of inhibitory pump cells (pump) and excitatory (Ine-NTS) interneurons in the NTS compartment.

The pontine IE population is included to intermingle the sensory input from the muscle afferent to brainstem respiratory populations. In addition, this population reciprocally interacts with medullary populations: it receives excitatory inputs from the post-I(e) and ramp-I populations providing inspiratory-expiratory modulation of its activity and giving excitatory input to the expiratory aug-E population (see Fig. M1 and Rybak et al. 2004).

The late-E population in RTN/pFRG is considered a population of central chemoreceptors sensitive to the level of CO_2_ and activated during hypercapnia (for detail, see Molkov et al. 2010). This population consists of neurons containing *I*_NaP_ and having mutual excitatory interactions within the population. During simulated hypercapnia, this population is activated by an external tonic drive. The following connections between the late-E population and other neural populations are incorporated in the model: (i) excitatory connections from the late-E population to the excitatory pre-I/I population of pre-BötC, allowing entrainment of the pre-BötC oscillations by the late-E oscillations; (ii) inhibitory connections from the inhibitory inspiratory population (early-I(1) of pre-BötC) to the late-E to provide inhibition of late-E neurons during inspiration (Fig. M1); (iii) excitatory connections from the late-E to the abdominal motor output (AbN) via bulbospinal premotor interneurons (bse), located in the cVRG compartment; (iv) excitatory connections from the late-E to the inhibitory populations aug-E (of BötC) and bsi (of rVRG), which both inhibit the premotor ramp-I population (see Fig. M1 and Molkov et al 2010); and (v) inhibitory connections from the post-I population of BötC to the late-E population.

The NTS compartment \was not present in the earlier models or was only included to provide pulmonary feedback to the brainstem respiratory network via pump cells (the pump population in the current model). However, this compartment includes a population of excitatory neurons sensitive to the level of O_2_ and activated during hypoxia (REF?). To simulate the effect of hypoxia on respiratory activity we included the population of excitatory neurons, Ine-NTS, in the NTS compartment of our model (see Fig. M1). During simulated hypoxia, this population is excited by external tonic drive and provides excitatory inputs to the late-E population in the RTN/pFRG, the ramp-I population in the rVRG, and to the HgMN population.

The left and right sides of the model interact by commissural connections via V0 neurons incorporated in the pre-I/I and ramp-I populations (Wu et al., 2017). These neurons had both ipsilateral local and descending projections: V0 neurons from the pre-I/I population project to the ipsi– and contralateral pre-I/I neurons and the ipsilateral ramp-I population; V0 neurons from the ramp-I population projected to the ipsi– and contralateral PhMNs populations. The spinal compartments in the model include local spinal respiratory microcircuits (see Fig. M1). Simulated muscle afferent inputs (via sensory interneurons populations, InS) activate PhMNs, spinal interneurons and the IE population in PONS (Potts et al. 2005 and Fig. M1). According to our hypothesis, the supraspinal pathways for activation of PhMNs in the model are normally suppressed by local inhibitory neurons (Ini) receiving the brainstem inputs and can be activated by stimulation of somatic afferents. We also incorporated commissural interneurons (CINe) at the spinal level (Fig. M1) to study potential effect of contralateral afferent stimulation. The model output is presented by integrated activity of phrenic and hypoglossal motoneuron populations simulating activities of the phrenic (PN) and hypoglossal (XII) nerves, respectively, and by simulated integrated activities of vagus (VN) and abdominal (AbN) nerves.

#### 3.2.2. Model validation. Simulation the effect of hypercapnia and hypoxia

The present model generalized, combined, and extended several earlier models (Molkov et al., 2010; Potts et al., 2005; Rybak, Shevtsova, Paton, et al., 2004; Smith et al., 2009), and preserved the advantages of the preceding basic models. Similar to these models it is able to reproduce multiple experimental data including the generation of eupneic respiratory pattern (Smith et al., 2007); changing this pattern following vagotomy (Rybak, Shevtsova, Paton, et al., 2004), and the quantal acceleration of abdominal activity following progressive hypercapnia (Molkov et al., 2010). To provide additional validation of the model, we have simulated the effects of hypercapnia and hypoxia and compare the results with the corresponding experimental data on hypercapnia (Ghali & Marchenko, 2016) and hypoxia (Marchenko unpublished) obtained in rats *in vivo*.

Hypercapnia was simulated by application of an additional external drive to both late-E populations in the left and right RTNs. The results simulations and ther comparison to the corresponding experimental results are shown in Fig. M2B1-B4. In both experimental and modeling results, hypercapnia produced an increase in hypoglossal activity. Both in the experiment and the model, hypoglossal bursts slightly preceded the phenic discharges in control conditions (see black arrows in Fig. M2B1,B3); however, during hypoglossal bursts started much earlier than the phrenic bursts (red arrows Fig. M3B2,B4). The period of the respiratory rhythm was not affected in both cases.

Hypoxia was simulated by simultaneous application of an additional external drive to Ine-NTS populations in the left and right NTS compartments (NTS receives inputs from peripheral chemoreceptors via the glossopharyngeal and vagus nerves, (Spyer & Gourine, 2009) and reduction by 20% of the tonic drive from the left and right pontine compartments to the medullary populations, see Table 3 in Appendix). The results of our simulation and comparison to the corresponding experimental results are shown in Fig. M2C1-C4. In the case of hypoxia, increase of hypoglossal activity becomes more pronounced in both experiment and simulation (red arrows in Fig. M2C2,C4) and the expiratory phase notably increases, resulting in slowing respiratory rhythm.

**Figure M2.**
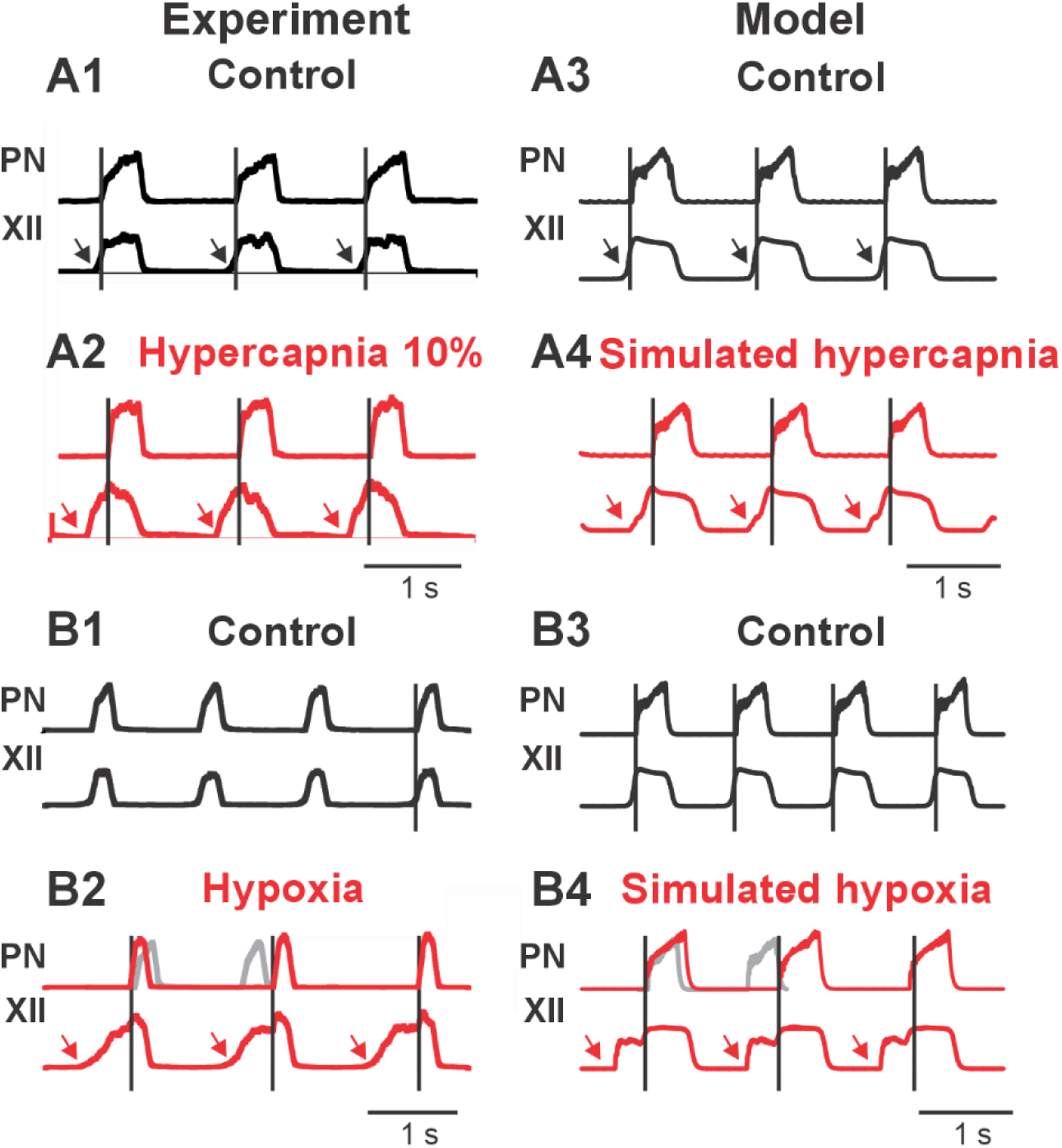
Comparison of the experimental and modeling results on rat respiration in challenging conditions. A1-A4: Response to hypercapnia in the experiment (A1,A2) and in the model (A3,A4). **B1-B4:** Response to hypoxia in the experiment (B1,B2) and in the model. Thick grey lines B2,B4 show where the next phrenic burst would occur without perturbation.

**Table 3.** Weights of synaptic connections in the network.

| Target population (location) | Excitatory drive (location)/weight of synaptic input or source population (location)/ mean weight of synaptic input from single neuron |
| --- | --- |
| IE (PONS) | post-I (e) (BötC)/2; ramp-I (rVRG)/2; InS(SC)/0.3; drive1 (PONS)/0.5 |
| pre-I /I (pre-BötC) | pre-I /I (pre-BötC)/0.3; c-pre-I /I (pre-BötC)/0.3; post-I (BötC)/-8; aug-E (BötC)/-0.1; late-E (RTN/pFRG)/2.12; drive1 (PONS)/0.75; drive2 (raphe)/0.75; drive3 (RTN)/0.13 |
| early-I (pre-BötC) | pre-I /I (pre-BötC)/0.85; c-pre-I /I (pre-BötC)/0.85; post-I (BötC)/-10; aug-E (BötC)/-0.1; pump (NTS)/-0.2; drive1 (PONS)/1.1; drive3 (RTN)/0.7 |
| post-I (BötC) | early-I (pre-BötC)/-2.4; aug-E (BötC)/-2; Ine-NTS (NTS)/0.8; drive1 (PONS)/1.45; drive3 (RTN)/0.05 |
| post-I (e) (BötC) | early-I (pre-BötC)/-4; aug-E (BötC)/-10; drive1 (PONS)/1 |
| aug-E (BötC) | early-I (pre-BötC)/-3.5; post-I (BötC)/4; late-E (RTN/pFRG)/0.9; IE (PONS)/1; drive1 (PONS)/0.2; drive3 (RTN)/1 |
| late-E (RTN/pFRG) | late-E (RTN/pFRG)/1.5; early-I (pre-BötC)/-0.8; post-I (BötC)/0.8; Ine-NTS (NTS)/1.38; hypercapnia/1 |
| ramp-I (rVRG) | pre-I /I (pre-BötC)/2; post-I (BötC)/-50; aug-E (BötC)/-10; bsi (rVRG)/-5; Ine-NTS (NTS)/1; drive1 (PONS)/1.9 |
| bsi (rVRG) | late-E (RTN/pFRG)/0.0; post-I (BötC)/-15; aug-E (BötC)/-2; drive1 (PONS)/3 |
| bse (cVRG) | late-E (RTN/pFRG)/3; bsi (rVRG)/-20; |
| Ine-NTS (NTS) | Hypoxia/1; Lungs/2 |
| pump (NTS) | Hypoxia/1; Lungs/2 |
| HgMN | pre-I /I (pre-BötC)/0.3 |
| Ini (SC) | ramp-I (rVRG)/2.5; c-ramp-I (rVRG)/2.5; InS(SC)/1; |
| Ine (SC) | ramp-I (rVRG)/2; c-ramp-I (rVRG)/2; Ini (SC)/; InS(SC)/0.5; CIne (SC)/1 |
| CIne (SC) | Ine (SC)/5 |
| RhMN (SC) | ramp-I (rVRG)/2; c-ramp-I (rVRG)/2; RC (SC)/-0.5; Ine (SC)/1; CIne (SC)/1 |
| RC (SC) | RhMN (SC)/1 |
| InS (SC) | Muscle afferents/1 |

\* Values after the “/” sign represent average weights of synaptic inputs from the corresponding drive ( $w_{dmi}$ ) or source populations, type ( $w_{ji}$ ), see Eq. (6). The “-” sign before connection weight indicates inhibitory connection. To provide heterogeneity of neurons within neural populations, the weights of excitatory and inhibitory connections were randomly distributed by $\pm 5\%$ or $\pm 10\%$ , respectively. Prefix c indicates contralateral population. All other connections are ipsilateral.

#### 3.2.3. Modeling results

##### Simulation of effects of afferent stimulation in intact rats

To reproduce our experimental results we first simulated the effects of biceps stimulation observed in our experiments in intact rats (Fig. M2A1-A4). Stimulation of biceps was simulated by introducing external drive to the InS population in the network (see Fig. M1) which excited the Ie population in the PONS and the aug-E population via pontine input to this population. As a result, application of a short external stimulation to the InS population at the end of the expiratory phase enhanced inhibition of the post-I population by the aug-E population in the BotC, thus producing premature switching to inspiration (Fig. M2 A4). Similar to the experimental results (Fig. 3B and M2A2), periodic sensory stimulation produced entrainment – premature termination of expiration and switching to inspiration in response to each stimulus (Fig. M2A4), Interestingly, in our simulation, unilateral stimulation produced the symmetric bilateral effect.

**Figure M3.**
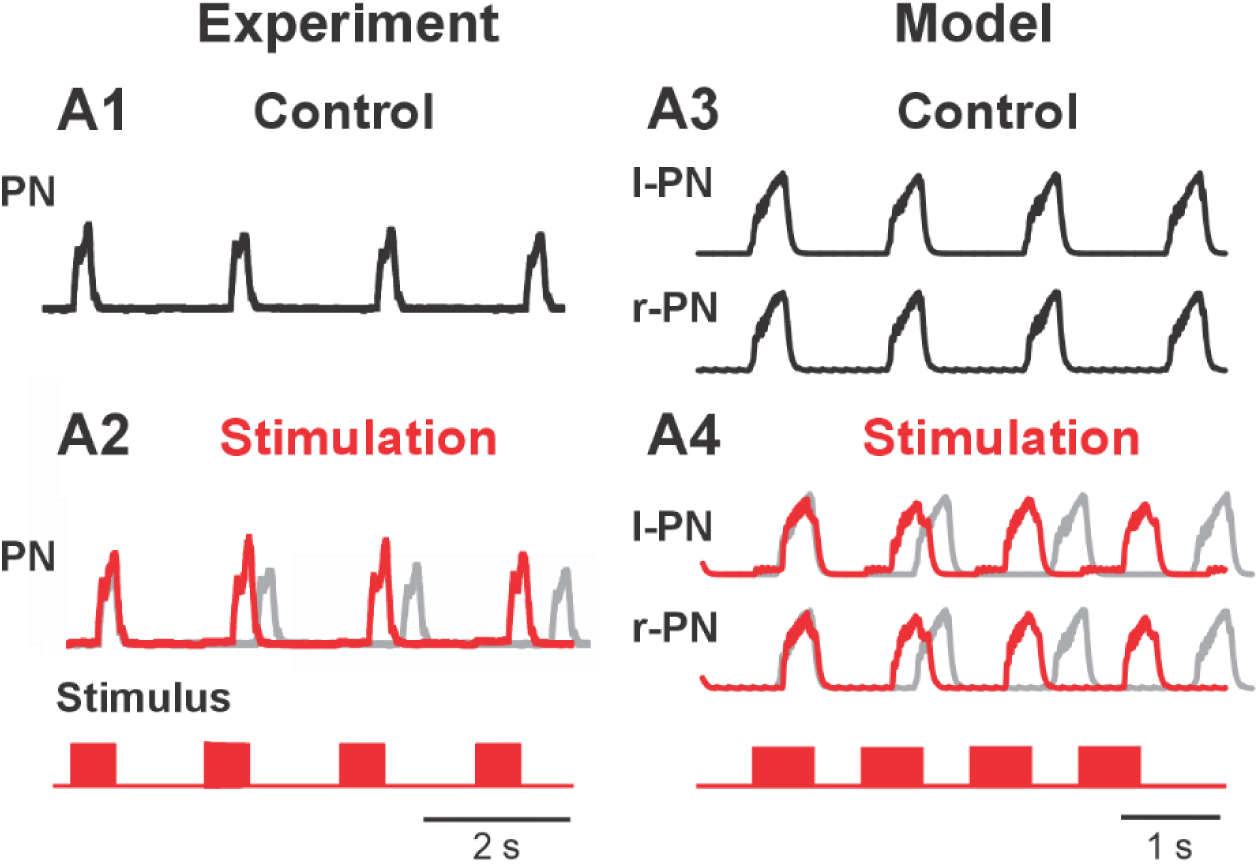
Simulation of effects of biceps stimulation in intact rats: A1, A2: Response to sensory stimulation in the experiment. A3, A4: Simulated response to afferent stimulation and in the model. Thick grey lines in A2 and A4 show where the next phrenic burst would occur without perturbation.

##### Simulation of effects of biceps stimulation in C2Hx rats

Based on our previously published work (Bezdudnaya, Hormigo, et al., 2018), we documented an acute spontaneous recovery of phrenic nerve activity immediately after injury. This early restoration suggests the presence of intrinsic compensatory mechanisms within the spinal and supraspinal respiratory networks that help reestablish functional output despite disrupted descending inputs. To further investigate the underlying dynamics of this early recovery, we simulated a spontaneous recovery of phrenic nerve activity using our computational model. Our simulations replicated these experimental findings and explored how circuit-level interactions could account for the observed restoration of phrenic motor activity.

Figure M4 illustrates the spontaneous recovery after hemisection at C2 level in rats and our simulation of this effect in the model. Hemisection was simulated by eliminating inputs from the medullary populations (light yellow boxes in Fig. M4A1,B1,C1) to the spinal compartment (light green boxes in Fig. M4A1,B1,C1) at the left side of the model. As shown in Fig. M4B2, this resulted in elimination of the ipsilesional phrenic activity while preserving contralesional rhythmic activity. To simulate the spontaneous recovery, the weights of connections from neurons of the ramp-I population contralateral to hemisection were increased by 50% which led to appearance of low amplitude bursts in the ipsilesional PN and increasing amplitude of the contralesional PN similar to experimental results (see Fig. M4C2,C3).

**Figure M4.**
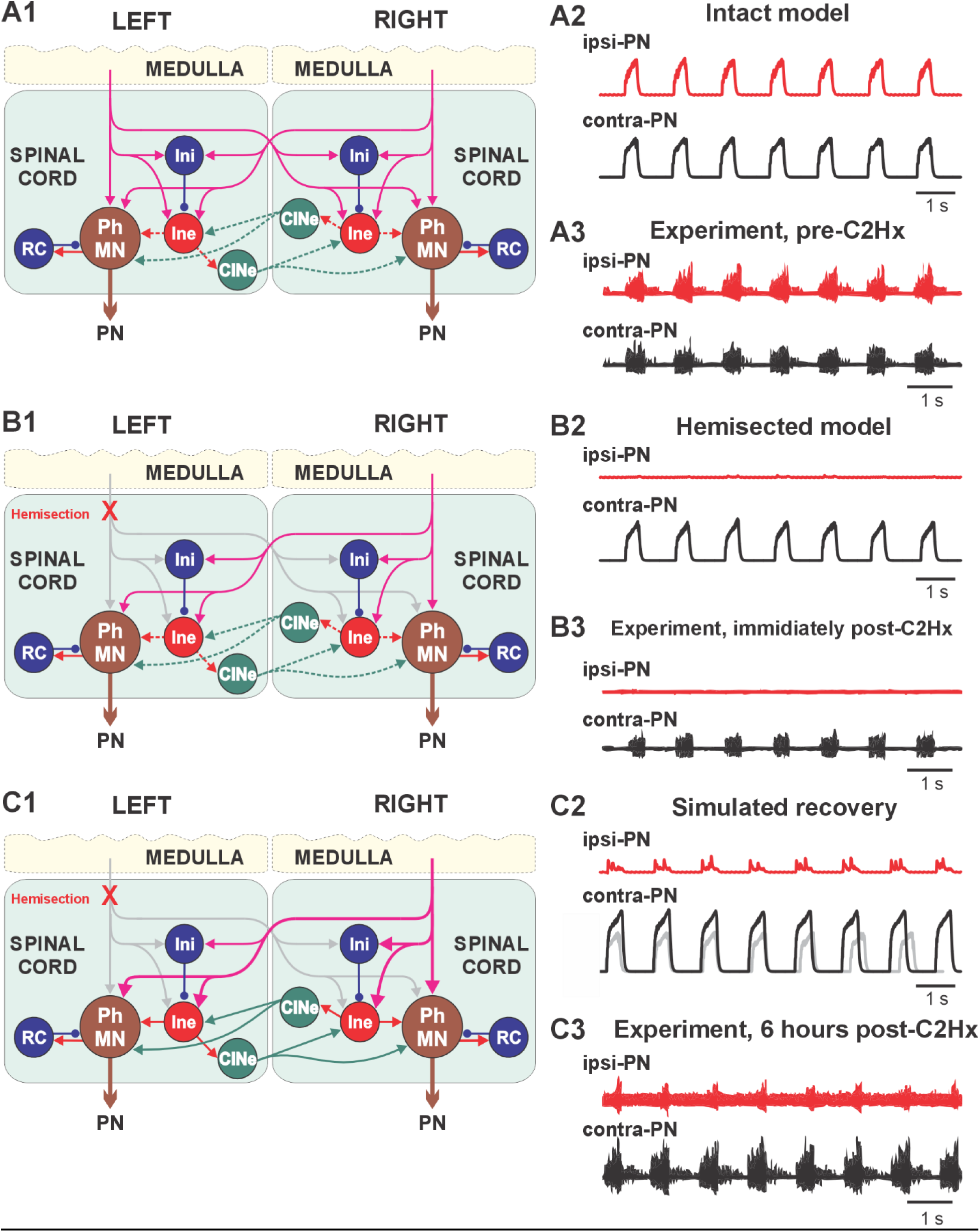
Simulation of spontaneous recovery of phrenic activity (PN) after hemisection at the C2 level (C2Hx). (A1,B1,C1) – The schematic diagrams illustrate changes in connectivity in the spinal respiratory circuitry following C2Hx. **(A2,B2,C2) –** Model performance in the intact case (A2), acutely following hemisection (B2), and after 6 h recovery (C2). In C2, thick grey trace shows the contra-PN activity in the intact model. Note the increase in amplitude of the contra-PN bursts after recovery. **(A3,B3,C3)-Experimental data:** Recovery of phrenic activity post-C2Hx in the rat. Phrenic motor outputs were determined by electrophysiological recording from the ipsilateral and contralateral to the injury phrenic nerves: ipsi-PN, contra-PN before, immediately, and 6 hours after C2Hx

Our next step was to reproduce and explain the experimental results (see Section 3.1.2) showing that biceps stimulation after C2 hemisection in rats can significantly increase the amplitude of spontaneous phrenic bursts on the injured side. To explore the mechanisms of this increase, we simulated biceps stimulation after hemisection. Figure M5 shows the results of our simulations. As in the intact case, stimulation was performed by inducing periodical additional excitatory drive to the InS population on the ipsilesional (left) side of the model. This drive directly increased activity of the left Ine population, and hence the activity of the ipsilesional PhMN population, and indirectly, via the commissural CINe populations, enhanced activity of both ipsi– and contralesional PhMNs. Modeling results demonstrated significant changes in ipsi-but not contralateral phrenic activity (ipsi-PN and contra-PN) resulting from sensory stimulation following hemisection similar to the experimental results (Fig. M5 C2 and D2, Fig. 4).

**Figure M5.**
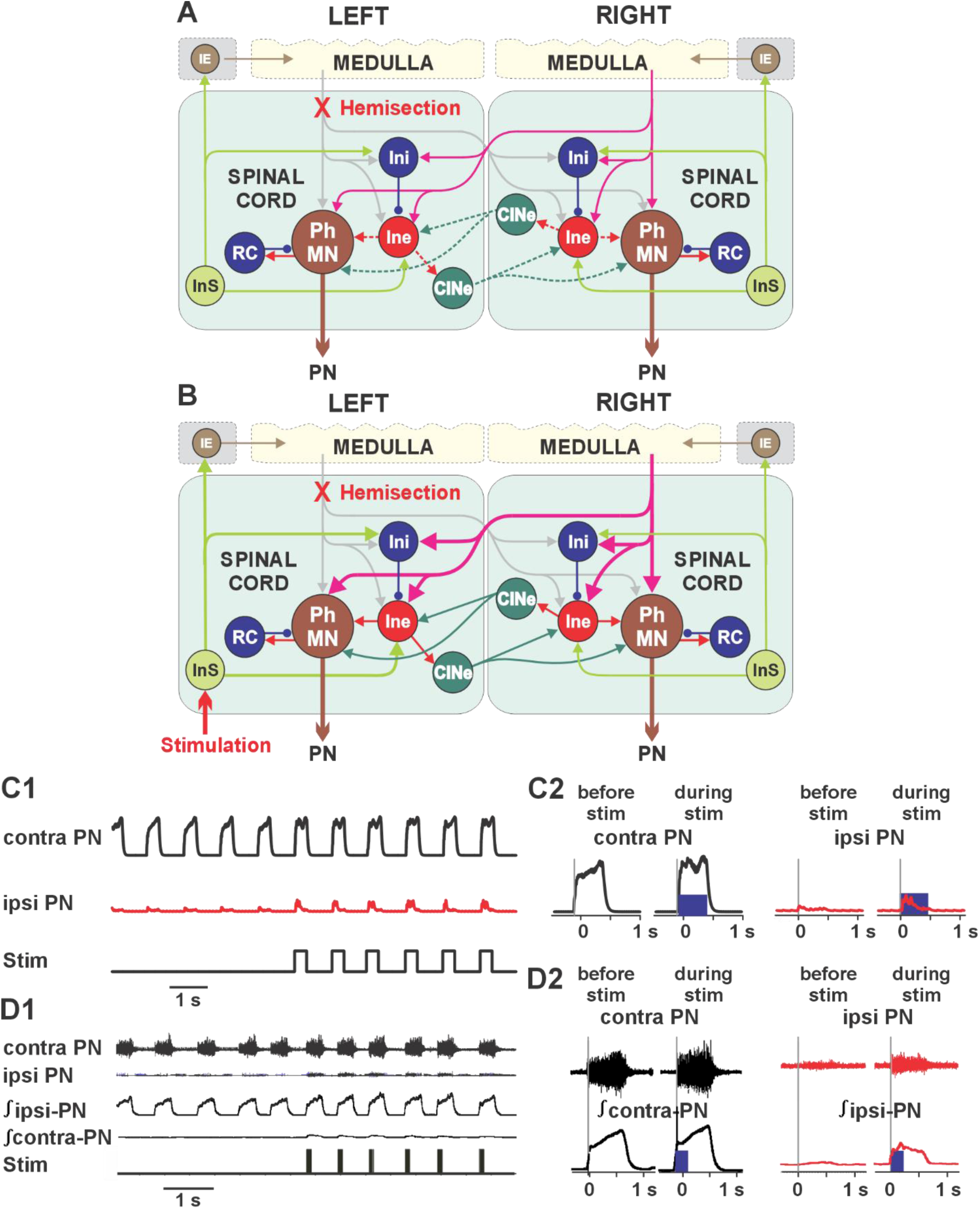
Simulation of the effect of biceps stimulation after hemisection at the C2 level. (A,B)
-The schematic diagrams illustrate connectivity in the spinal respiratory circuitry following hemisection (A) and its change after recovery followed by simulated sensory stimulation (B). **(C1, C2)-** Modeling results demonstrate changes in ipsi– and contralateral phrenic activity (Ipsi-PN and contra-PN) resulting from sensory stimulation following hemisection. (D1,D2)-experimental data used for model simulation.

##### Simulation of effects of biceps stimulation in C1Tx rats

C1 transection eliminates all descending inputs from the brainstem to the respiratory spinal circuits, resulting in a total loss of phrenic nerve activity. It might be suggested that electrical stimulation of the biceps could evoke a phrenic activity as in the case of C2Hx. However, it was shown that afferent stimulation after C1Tx had no effect on phrenic nerve activity. We have hypothesized that the effect of afferent stimulation in the case of C1Tx is suppressed by inhibitory influences of some spinal interneurons and this suppression could be eliminated by blockade of inhibitory synaptic connections. To verify this hypothesis, we simulated the effect of complete transection at the C1 level by eliminating all supraspinal inputs to the spinal compartment and blocked the inhibition of the Ine populations by Ini populations on both sides of the model (Fig. M6 A and B). Figure M6C shows the results of our simulations and demonstrates that disinhibition of both left and right Ine populations and the periodical excitation of the left Ine population by sensory stimulation triggered activity in the left (ipsilateral to stimulation) PhMn population and, via the commissural CINe populations and the right Ine population, in the right (contralateral to stimulation) PhMN population. Our prediction was further confirmed in the experiments (see section 3.1.3 and Fig. 5).

**Figure M6.**
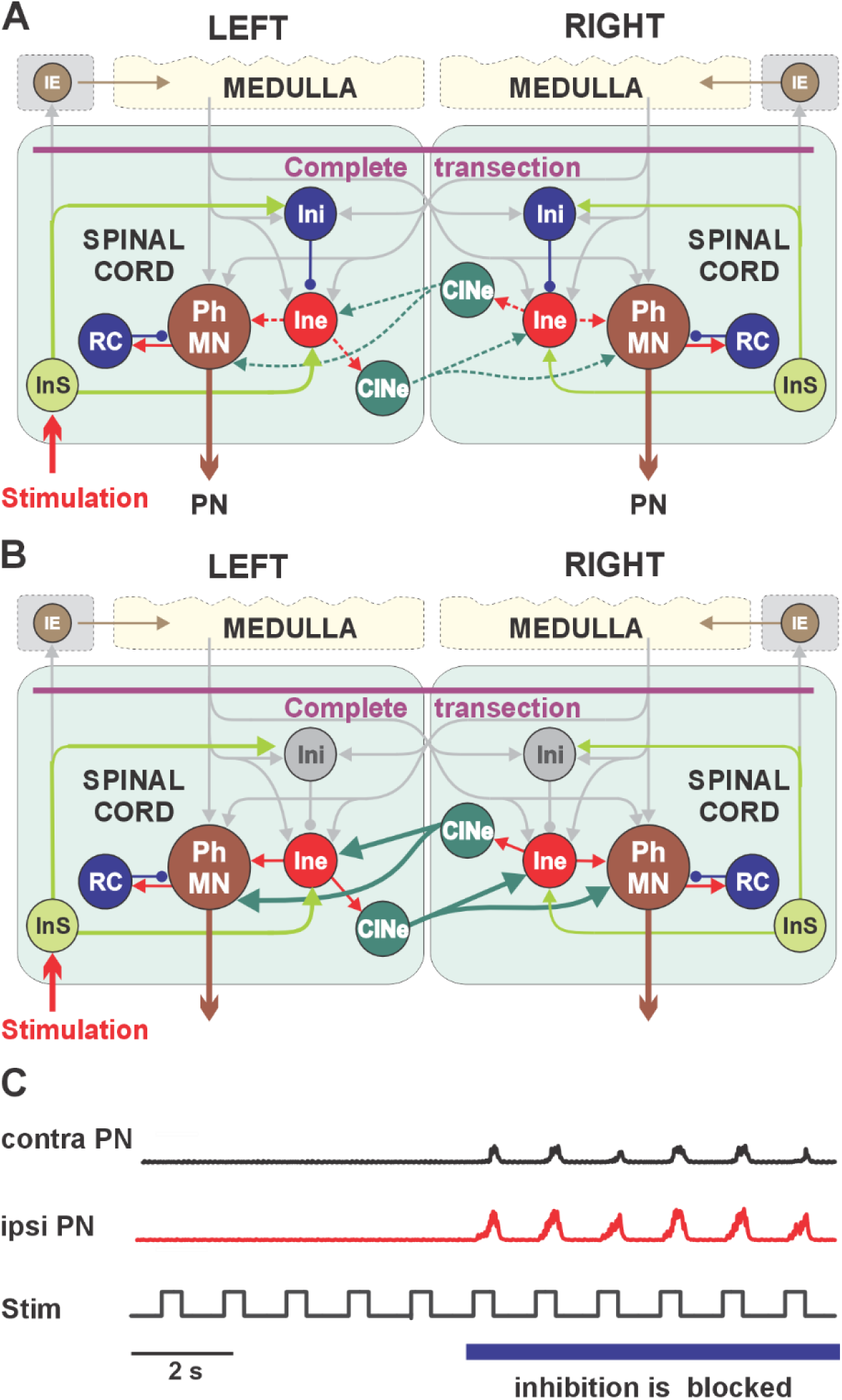
Simulation of the effect of biceps stimulation in C1 transected rat when blocking synaptic inhibition in the spinal compartment. (A,B) The schematic diagrams illustrate connectivity in the spinal respiratory circuitry following complete transection spinal (A) and its change after simulated sensory stimulation when synaptic inhibition in the spinal compartment in the model was blocked (B). **(C1, C2)** Modeling results demonstrating ipsi– and contralateral to the stimulation phrenic activity (Ipsi-PN and contra-PN) in the transected model during sensory stimulation before and after blockade of synaptic inhibitory transmission in the spinal compartment.

## 4. Discussion

In this paper, we demonstrate how forelimb muscle stimulation affects phrenic nerve activity and respiration in spinal cord intact, C2-hemisected, and C1-transected rats, and present a comprehensive computational model of respiratory circuits that simulates experimental data and provides viable predictions.

While both supraspinal and spinal mechanisms contribute to the regulation of phrenic motor neuron (PhMN) activity and breathing, our primary interest lies in the contributions of spinal-level interactions. We suggest that the spinal cord contains integrative networks that coordinate PhMN activity with other motor behaviors and sensory inputs.

According to our model (Fig. 6), the following mechanisms can be hypothesized:

1. Spinal integration: Circuits controlling PhMNs interact with other motor networks within the spinal cord, enabling adaptive modulation of breathing during movement and other behaviors.
2. Local premotor control: The activity of PhMNs on each side of the spinal cord is shaped primarily by local excitatory and inhibitory premotor interneurons that relay descending respiratory drive from the brainstem.
3. Sensory modulation: Afferent inputs from limb muscles influence these premotor interneurons through ipsilateral and commissural spinal pathways, allowing sensory feedback to modulate respiratory output.
4. State-dependent recruitment: Under normal conditions, these intraspinal pathways are tonically inhibited by local interneurons but can be recruited during specific physiological states such as locomotion, exercise, or respiratory challenges.

**Figure. 6.**
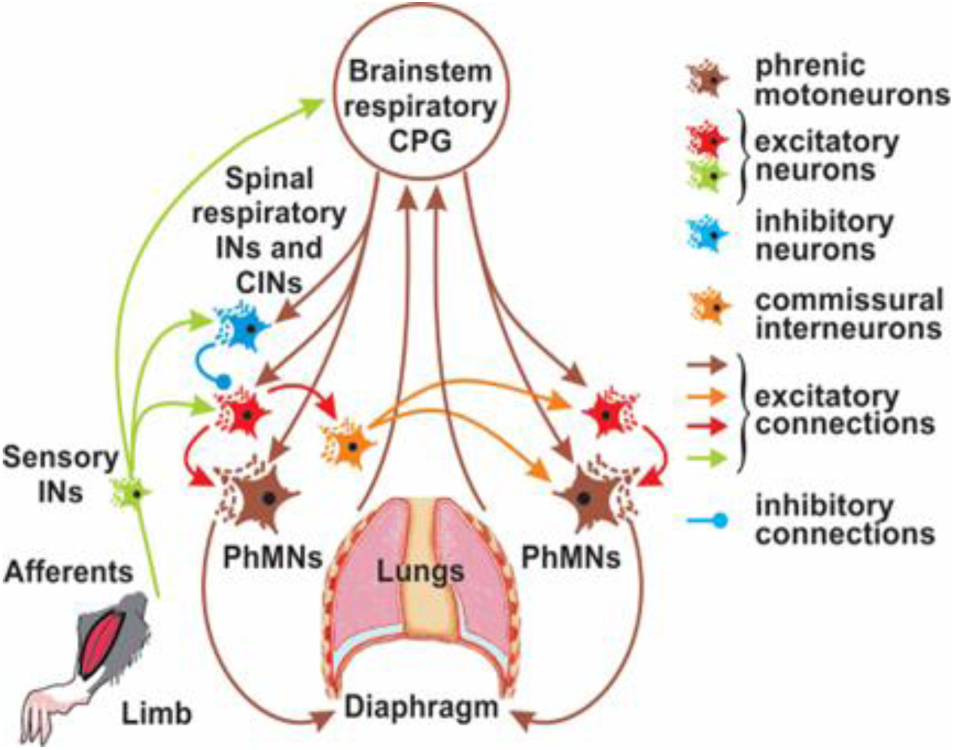
Conceptual model of spinal circuits coordinating respiratory and limb motor networks.

Activation of considered spinal pathways may represent a mechanism for enhancing or restoring respiratory function after spinal cord injury.

### 4.1. Modulation of respiratory output by upper limb muscle stimulation in intact animals

In spinally intact rats, biceps muscle stimulation produced rapid increases in both tidal volume and breathing frequency (Fig. 2), indicating that forelimb afferent input can modulate respiratory rhythm generation (Giraudin et al., 2012; Le Gal et al., 2020; Morin & Viala, 2002; Potts et al., 2005). Nevertheless, the short-lived nature of these responses suggests that supraspinal mechanisms rapidly re-establish the intrinsic respiratory rhythm, maintaining homeostatic control and limiting sustained influence from peripheral inputs. In chemo– and baro-denervated, mechanically ventilated rats, changes in phrenic activity were more difficult to elicit, likely because the removal of peripheral feedback reduces the overall excitability of the respiratory system, making it more stable. Interestingly, in a subset of these experiments, rhythmic biceps stimulation could still entrain phrenic activity, indicating that even in the absence of classical sensory inputs, somatic feedback can synchronize respiratory motor output through spinal–brainstem circuits (Fig. 3).

Interestingly, studies in mice using sciatic nerve electrical stimulation also reported a transient effect of stimulation on diaphragm activity in intact animals (Walling et al., 2024). However, in our data, we did not observe an increase in tonic phrenic nerve activity during stimulation. It is important to note that stimulation protocol of the work above differed from ours, that work applied continuous 20 Hz stimulation for 1 minute, whereas we used a burst stimulation paradigm (see Methods). We believe that burst stimulation produces a clearer and more physiologically relevant effect.

### 4.2. Enhancement of phrenic output by biceps stimulation following cervical hemisection

In animals with a C2 hemisection, stimulation of the ipsilateral biceps brachii significantly increased phrenic nerve amplitude on the injured side (Fig. 4). This effect is consistent with previous findings in mice, where sciatic nerve stimulation also enhanced diaphragm activity after C2Hx (Walling et al., 2024). Overall, these results indicate that forelimb afferent feedback can increase respiratory motoneuron excitability and engage residual spinal circuits even after partial loss of descending input. The absence of a significant contralateral effect likely reflects ongoing supraspinal control of the intact side and compensation for the loss of diaphragmatic activity ipsilaterally (Bezdudnaya, Hormigo, et al., 2018; Bezdudnaya et al., 2017; Lee & Hsu, 2017; Sandhu et al., 2009). As a result, contralateral phrenic output remains more stable and less influenced by peripheral stimulation. Overall, these findings align with the concept of locomotor–respiratory coupling, where sensory input from limb movement modulates respiratory output. Following spinal cord injury, spinal mechanisms may become increasingly important as supraspinal connectivity diminishes, providing an alternative pathway for respiratory modulation and functional recovery.

The computational model reproduces two key phenomena observed experimentally following C2 hemisection: the acute spontaneous recovery of ipsilesional phrenic activity (Fig. M4) and its further enhancement by forelimb afferent stimulation (Fig. M5). The early re-emergence of low-amplitude ipsilesional phrenic bursts in the simulations supports the idea that intrinsic network rebalancing, particularly the strengthened influence of contralateral inspiratory drive, can partially compensate for the loss of direct descending inputs. Biceps stimulation amplifies this recovered activity by directly increasing the excitatory drive to ipsilesional phrenic motor neurons through local premotor interneurons and indirectly enhancing contralateral phrenic activity via commissural interneurons. However, this indirect enhancing of contralateral phrenic motoneurons were not enough to elicit significant changes similar to our experimental data (Fig. 4). These findings suggest that early post-injury respiratory recovery arises from dynamic interactions within the pre-existing bilateral network and sensory-driven spinal mechanisms, rather than from structural reorganization, and highlight the capacity of somatic afferent input to enhance motor output by leveraging residual circuit interactions. An important question is whether our biceps stimulation produces pain and alters respiratory rate or strength. Most of the rats used in these experiments are decerebrate, so they cannot process pain at the cortical level; however, sympathetic reflexes remain intact. One of the indicators of pain is changes in blood pressure (Sacco et al., 2013). It may increase during painful stimulation but in our experiments, it could change transiently but typically return quickly to baseline. We also monitor pupil size during stimulation. Pain would cause pupillary dilation (mydriasis) (Wildemeersch et al., 2018), which we did not observe. Lastly, our stimulation parameters were within the normal physiological range, and we believe they did not activate C or D pain fibers.

### 4.3. Respiratory responses after complete transection under pharmacological disinhibition

Prior to conducting these experiments, we used the developed computational model to simulate the effects of cervical spinal cord transection, pharmacological disinhibition, and both unilateral and bilateral biceps stimulation (Fig. M6). Remarkably, all experimental outcomes predicted by the model were subsequently confirmed in vivo experiments, providing strong validation for the model’s accuracy and predictive power. First, biceps stimulation failed to evoke any phrenic activity after complete C1 transection, confirming that under normal inhibitory conditions, forelimb afferent input alone is insufficient to activate phrenic motoneurons without descending drive. However, following pharmacological disinhibition with Gabazine and Strychnine, unilateral muscle stimulation produced robust phrenic bursts. Consistent with the model predictions (Fig.M6), the responses were bilateral, with larger and faster activation on the ipsilateral side, indicating that latent excitatory pathways between limb afferents and respiratory circuits remained intact below the lesion and can be unmasked when inhibitory tone is reduced. Bilateral muscle stimulation produces approximately same latencies and amplitudes responses on both sides (Fig. 5).

The question is whether diaphragm contractions elicited by biceps stimulation in C1Tx animals are sufficient to support vital respiration. Our preliminary data suggest that this may be possible. Figure 7 shows recordings from a spontaneously breathing, C1Tx decerebrate rat before and after the ventilator was turned off and on, with tracheal pressure and airflow recorded instead of phrenic nerve activity. Gabazine and strychnine were applied prior to the experiment. When the ventilator was switched off, biceps stimulation produced negative airflow, indicative of inspiration. Even though stimulation was delivered at a low frequency of one burst every 5 seconds, the animal was able to maintain breathing under stimulation for approximately 10 minutes, after which tracheal pressure began to decline and end-tidal CO₂ increased, prompting re-initiation of mechanical ventilation. These results indicate that, although diaphragm activation via biceps stimulation can generate inspiratory airflow for a limited period, further experiments are required to optimize stimulation parameters.

**Figure 7.**
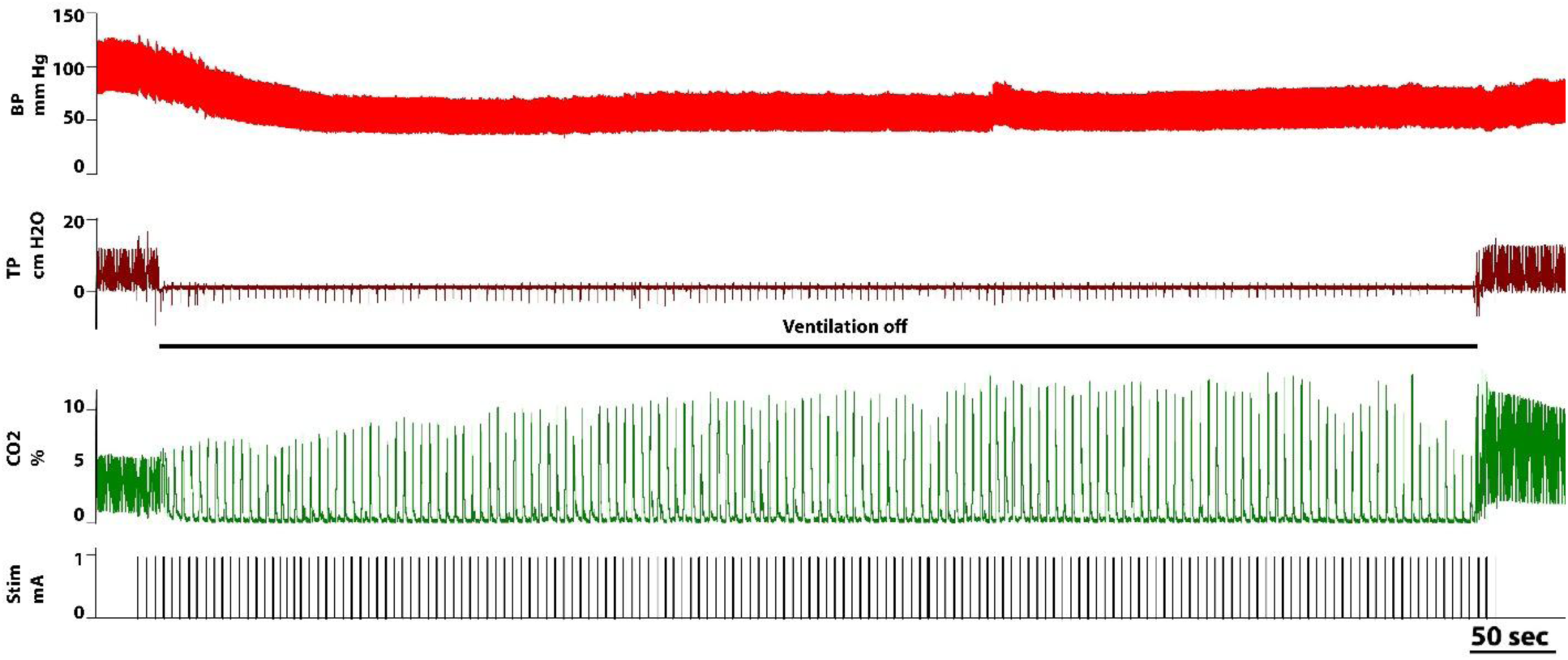
Example of biceps stimulation in T1Tx rat during ventilation and without stimulation. Abbreviations: BP-blood pressure, TP –tracheal pressure, CO2 –end tidal CO2, Stim. – biceps stimulation.

Interestingly, previous studies have shown that spinal cord injury disrupts the balance between excitatory and inhibitory signaling below the lesion, often leading to reduced inhibition. A valuable future direction would be to examine how biceps stimulation affects phrenic activity during the chronic phase of injury, when altered inhibitory control, such as downregulation of KCC2 (Cheung et al., 2023; Talifu et al., 2022), contributes to motor circuit hyperexcitability. Notably, sciatic nerve stimulation in chronically injured C2Hx mice has also produced better results than in acutely injured animals (Walling et al., 2024). Elucidating these mechanisms could provide important insight into how limb stimulation modulates respiratory output and help develop strategies to restore functional balance within spinal networks.

## 5. Conclusions

In summary, this study provides converging experimental and computational evidence for functional coupling between forelimb motor circuits and respiratory networks within the cervical spinal cord. Our findings demonstrate that limb afferent input can modulate phrenic motor output through intraspinal mechanisms that depend on descending drive, local inhibitory tone, and preserved interneuronal connectivity. Following cervical spinal injury, these spinal interactions can partially compensate for disrupted supraspinal control, and under conditions of reduced inhibition, previously latent pathways can be recruited to enhance respiratory motor output. Together, these results support a model in which spinal motor circuits dynamically interact to adapt breathing to motor demands and identify a spinal substrate that may be targeted to promote respiratory recovery after cervical spinal cord injury (Shevtsova et al., 2019).

## Author contributions

NS: computational modeling, data interpretation and manuscript writing; VM: performed experiments and data analysis, manuscript writing; IR: computational modeling and manuscript writing; TB: conception and design of the work, assistance with experiments and data analysis, data interpretation, and manuscript writing. All authors (NS, VM, IR, TB) approved the final version of the manuscript.

## Appendix

### Modeling formalism and parameters

All neurons were modelled in the Hodgkin-Huxley style as single-compartment models. The dynamics of their membrane potentials was described by the following differential equation:

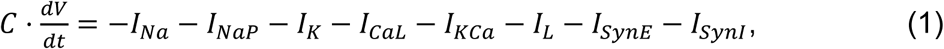

where *V* is the neuron membrane potential, *C* is the membrane capacitance, and *t* is time. The terms in the right part of this equation represent ionic and synaptic currents included in the neuron model: *I_Na_* – fast sodium (with maximal conductance ̄*g*_*Na*_); *I_NaP_* – persistent (slow inactivating) sodium (with maximal conductance ̄*g*_*NaP*_); *I_K_* – delayed-rectifier potassium (with maximal conductance ̄*g*_*K*_); *I_CaL_* – high-voltage activated calcium-L (with maximal conductance ̄*g*_*CaL*_); *I_KCa_* – calcium-dependent potassium (with maximal conductance ̄*g*_*K*,*Ca*_), *I_L_* – leakage (with constant conductance *g_L_*); *I_SynE_* – excitatory synaptic (with conductance *g_SynE_*), and *I_SynI_*–inhibitory synaptic currents (with conductances *g_SynI_*).

In Eq. (1), the ionic and synaptic currents are described as follows:

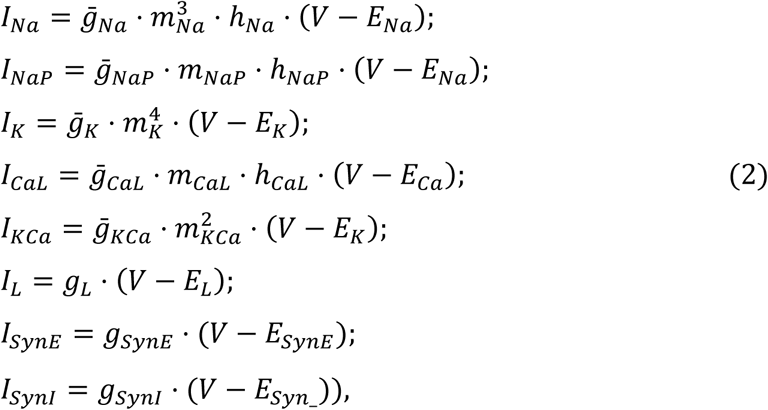

where *E_Na_*, *E_K_*, *E_Ca_*, *E_L_*, *E_SynE_*, and *E_SynI_* are the reversal potentials for the corresponding membrane and synaptic channels.

Variables *m_i_* and *h_i_* with indexes indicating ionic currents represent, respectively, the activation and inactivation variables of the corresponding ionic channels. Kinetics of activation and inactivation variables for all voltage–dependent currents are described as follows:

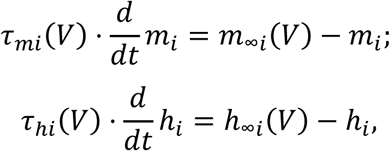

where *m_∞_*_*i*_and ℎ*_∞_*_*i*_ are steady states for activation and inactivation variables, respectively, τ_*mi*_ and τ_ℎ*i*_ are the corresponding time constants. The expressions for steady state activation and inactivation variables and time constants are shown in Table 1. Activation of calcium-dependent potassium channels (*KCa*) depends on the intracellular calcium concentration and is independent of voltage (see Table 1).

The kinetics of intracellular Ca^2+^ concentration, *Ca*, is modeled according to the following equation (Booth et al. 1997):

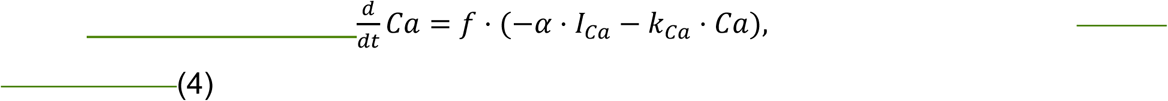

where *f* defines the percent of free to total Ca^2+^; α converts the total Ca^2+^ current, *I_Ca_*, to Ca^2+^ concentration; *k_Ca_* represents the Ca^2+^ removal rate. In the current study, *I_Ca_* is represented by the high-voltage activated calcium-L current, *I_CaL_* (see Eqs. (1) and (2) and Table 1).

Activation of the Ca^2+^-dependent potassium channels is also considered instantaneous and described as follows (Booth et al. 1997):

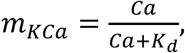

where *Ca* is the Ca^2+^ concentration within the corresponding compartment or neuron, and *K_d_* defines the half-saturation level of this conductance.

The maximal conductance values for all neuron types are shown in Table 2.

The excitatory (*g_SynE_*) and the inhibitory (*g_SynI_*) synaptic conductances are equal to zero at rest and may be activated (opened) by the corresponding synaptic input:

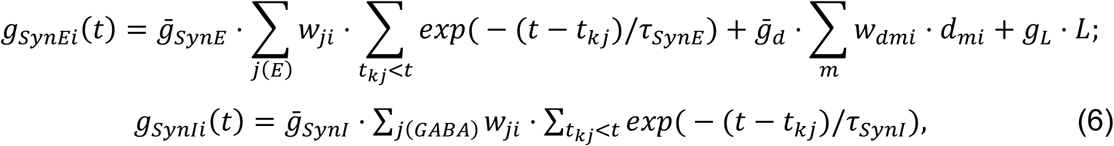

In Eq. (6), each type of synaptic conductance for neuron *i* (the excitatory, *g_SynEi_(t)*, and inhibitory, *g_SynIi_(t)* are summarized over all inputs *j* of the corresponding type (indicated under the first sigma sine). The first equation in (6) for the excitatory synaptic conductance has a second term describing the integrated effect of inputs from external drives, *d_mi_*. Each spike arriving to excitatory or inhibitory input of neuron *i* from neuron *j* at time *t_kj_* increases the corresponding synaptic conductance by ̄*g*_*SynE*_ ⋅ *w*_*ji*_, or ̄*g*_*SynI*_ ⋅ *w*_*ji*_, respectively, where ̄*g*_*SynE*_ and ̄*g*_*SynI*_ are the parameters defining an increase in the corresponding synaptic conductance produced by one arriving spike at synaptic weight *w_hy_* = 1. τ_*SynE*_ and τ_*SynI*_ are the decay time constants for the corresponding conductances respectively. In the second term of the first equations, ̄*g*_*d*_ is the parameters defining the increase in the excitatory synaptic conductance, respectively, produced by external input drive *d_mi_* = 1 with a synaptic weight of *d_im_* = 1. All drives were set equal to 1. The average weights of synaptic connections (*w_ij_* and *d_im_*) are shown in Table 2, Appendix 1. In the third term, *g*_*L*·_ = 1 is the connection weight from pulmonary stretch receptors to the pump population in the NTS (see Fig. M1) and *L* is the integrated activity of the pulmonary stretch receptors providing the feedback from the diaphragm to the respiratory center in the medulla. The simplified models of lungs and slowly adapting pulmonary stretch receptors is described as follows:

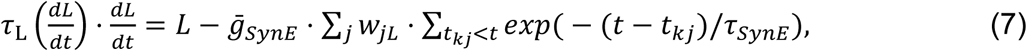

where

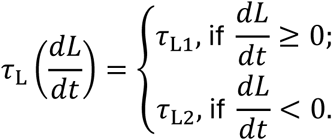

The second term in the right part of Eq. (7) describes the integrated effect of the excitatory inputs from the left and right PhMN populations to the diaphragm. Each spike arriving to the diaphragm from neuron *i* of the left and right PhMN populations at time *t_kj_* increases the corresponding synaptic input by ̄*g*_*SynE*_ ⋅ *w*_*ji*_, where ̄*g*_*SynE*_ is the parameters defining an increase in the excitatory synaptic conductance produced by one arriving spike at synaptic weight *w_hy_* = 1. τ_*SynE*_ are the decay time constant for the excitatory conductance.

The following general model parameters were used: capacitance: C = 36 pF; synaptic parameters: ̄*g*_*SynE*_= ̄*g*_*synI*_=̄*g*_*d*_ = 1.0 nS; *τ_SynE_* = 5 ms, *τ_SynE_* = 15 ms; parameters of calcium kinetics: *f* = 0.01; α = 0.0009 mol/(C×µm); *k_Ca_* = 2 ms^−1^; *K_d_* = 0.2 µM; reversal potentials: *E_Na_* = 55 mV; *E_K_* = –94 mV; *E_SynE_* = –10 mV; *E_SynI__* = –75 mV; lung parameters: τ_L1_ = 20; τ_L2_ = 200; *w*_*jL*_ = 1.

